# Compact Tape-Driven Sample Delivery System for Serial Femtosecond Crystallography

**DOI:** 10.1101/2025.09.25.673893

**Authors:** Jungmin Kang, Yoshiaki Shimazu, Fangjia Luo, Ayumi Yamashita, Tomoyuki Tanaka, Yuichi Inubushi, Kensuke Tono, Nipawan Nuemket, Allen M. Orville, So Iwata, Eriko Nango, Makina Yabashi

**Affiliations:** RIKEN SPring-8 Center, 1-1-1, Kouto, Sayo-cho, Sayo-gun, Hyogo, 679-5148, Japan; Japan Synchrotron Radiation Research Institute, 1-1-1, Kouto, Sayo-cho, Sayo-gun, Hyogo, 679-5198, Japan; Diamond Light Source, Harwell Science and Innovation Campus, Didcot, Oxfordshire OX11 0DE, United Kingdom; Research Complex at Harwell, Harwell Science and Innovation Campus, Didcot, Oxfordshire OX11 0FA, United Kingdom; Kyoto University, Yoshida-konoe-cho, Sakyo-ku, Kyoto, 606-8501, Japan; Tohoku University, 2-1-1 Katahira, Aoba-ku, Sendai, 980-8577, Japan

**Keywords:** Serial femtosecond crystallography, X-ray free-electron laser, room-temperature crystallography, conveyor belt, drop-on-tape, nanoliter droplet

## Abstract

We developed a compact tape-drive with on-demand sample delivery (CoT) for time-resolved serial femtosecond crystallography (SFX) experiments that can deliver sample droplets and/or initiate reactions with a drop-on-drop strategy. Two disposable piezoelectric injectors are positioned in tandem along the tape to produce a queue of nanoliter-scale droplets. X-ray free-electron laser (XFEL) pulses arrive perpendicular to and pass through the broad face of the tape where the pulse is synchronized and aligned to the droplets and thereby enables very highly efficient SFX data collection. The tape transport speed and the delivery distance can be varied to control the mixing time from approximately 130 ms to tens of seconds. We conducted time-resolved SFX experiments utilizing a basic enzymatic reaction model of hen egg white lysozyme (HEWL) and *N*-acetyl-*D*-glucosamine (GlcNAc) to demonstrate the drop-on-drop capabilities of CoT, and the full binding process of GlcNAc to HEWL was observed at 1.3–9.7 s.

**Synopsis:** Tape-driven liquid sample droplet delivery system for serial femtosecond crystallography

## 1. Introduction

Serial femtosecond crystallography (SFX) is a method to determine crystal structures using an X-ray free-electron laser (XFEL) (Chapman *et al*., 2011, Barends *et al*., 2022). In SFX, diffraction patterns are collected from randomly oriented microcrystals under room temperature conditions before the onset of radiation damage with intense, femtosecond X-ray pulses (Neutze *et al*., 2000). When combined with reactions triggered by light, mixing with a substrate, and/or heating, these diffraction patterns measured at near-physiological temperatures allow for the real-time observation of the structural dynamics and chemical reactions that occur in protein crystals. Time-resolved SFX (TR-SFX), which combines a reaction initiator and SFX, is widely used to visualize the structural changes and reactions in proteins (Tenboer *et al*., 2014).

SFX requires pristine microcrystals to be supplied to the XFEL intersecting area for each pulse because the crystals are damaged after irradiation. Thus, various sample delivery methods have been employed to supply crystals to the XFEL intersecting area that can be classified into two main types: continuous flows and discrete droplets. The continuous type, such as liquid jet and high-viscosity sample injectors, carry crystals suspended in a buffer or embedded in a high-viscosity carrier. A typical SFX experiment requires a crystal slurry with a high crystal density (crystals mL^−1^) to achieve the optimal hit rate (i.e., percentage of XFEL pulses diffracted by crystals) of 20–60%. Thus, crystals are wasted during the interval between XFEL pulses, which increases sample consumption especially at low repetition rates of several tens of hertz. The discrete type includes our previously reported pulsed liquid droplet injector (Mafune *et al*., 2016), which ejects around 0.3 nL droplet with an inner-diameter of 80 µm nozzle containing microcrystals from a piezo-driven droplet nozzle in synchronization with an XFEL pulse, and acoustic droplet ejection (ADE) (Roessler *et al*., 2016), which uses focused sound waves to eject a 2–3 nL volume droplet into an XFEL pulse. Discrete-type sample delivery methods do not waste crystals during the interval between XFEL pulses and thus reduce the sample consumption. However, ensuring that XFEL pulses consistently hit crystals in micron-scale droplets is a major challenge especially if the crystals are heterogeneous in size, which affects the droplet ejection speed and stability. In addition, droplets move at approximately 14 m s^−1^, and their trajectory wanders more with longer travel distance, both of which limit the delay time for TR-SFX experiments to around 2 µs (Kubo *et al*., 2017).

To address these challenges, Fuller *et al*. introduced the drop-on-tape (DOT) method that utilizes a tape as a conveyor belt to form a sequence of droplets and ensure their precise arrival at the XFEL intersecting area (Fuller *et al*., 2017). Their setup uses ADE (Roessler *et al*., 2016) to eject 0.8–6.0 nL droplets synchronized to XFEL pulses on the tape, which is driven at a speed of 30–600 mm s^−1^. The XFEL is positioned parallel to the tape surface to enable simultaneous SFX and X-ray emission spectroscopy (XES) from each drop and X-ray pulse. After irradiation, the tape is cleaned and reused for subsequent droplets. To demonstrate the capabilities of the DOT method, Fuller *et al*. conducted pump-probe experiments using light excitation with photosystem II and time-resolved experiments involving the gas activation of ribonucleotide reductase R2. They also conducted time-resolved mixing experiments by combining the DOT method with an additional piezoelectric injector that dispensed picoliter-scale ligand droplets onto a nanoliter-scale sample droplet to induce turbulence for mixing as the droplets merged (Butryn *et al*., 2021). Butryn *et al*. used hen egg white lysozyme (HEWL) and *N*-acetyl-*D*-glucosamine (GlcNAc) and were able to visualize the process of GlcNAc binding to active sites over time; whereas Nguyen *et al*. used a CYP121 (a P450 enzyme from *Mycobacterium tuberculosis*) and peracetic acid binding to initiate a peroxide shunt reaction (Nguyen *et al*., 2020).

Here, we report a compact tape-driven sample delivery system (CoT) for mix-and-inject SFX experiments based on the DOT. Instead of positioning the XFEL parallel to the tape, CoT positions the XFEL so that the pulses penetrate a droplet on the tape vertically to facilitate alignment between the XFEL intersecting area and sample droplets. While the DOT method cleans and dries the tape *in situ* for continuous use over extended periods of time, CoT utilizes a reel-to-reel tape drive and single-use tapes. Piezoelectric injectors with disposable nozzles are used to eject sample droplets. To demonstrate the capabilities of CoT, we conducted time-resolved mixing SFX experiments at the SPring-8 Angstrom Compact free-electron laser (SACLA) facility using HEWL crystals and GlcNAc.

## 2. Compact tape-driven sample delivery (CoT) setup

CoT comprises a drive wheel, tape reels, crown rollers, disposable piezoelectric injectors with sample supply components, a lift unit for changing the distance between the samples and X-ray interaction point, and stepping motors, which are all mounted on a 12-mm-thick duralumin plate. Figure 1a shows CoT mounted on the Diverse Application Platform for Hard X-ray diffractioN In SACLA (DAPHNIS) (Tono *et al*., 2015), which is equipped with a multiport charge-coupled device (MPCCD) detector (Kameshima *et al*., 2014). The main body is 400 mm wide and 730 mm high, and the *x*-, *y*-, and *z*-axis stages for aligning the sample droplets with the XFEL intersecting area are installed below the body. A collimator through which XFEL beams pass is fixed to the DAPHNIS stand and is separated from the main body plate. Helium gas flows inside the collimator to prevent parasitic scattering while the entire setup is kept under atmospheric conditions. Cameras are used to monitor the ejection of the sample droplets and their delivery to the XFEL intersecting area. This device is also designed to enable pump-probe TR-SFX experiments. An optical system for photoexcitation of crystals on the tape can be introduced into the setup.

**Figure 1.**
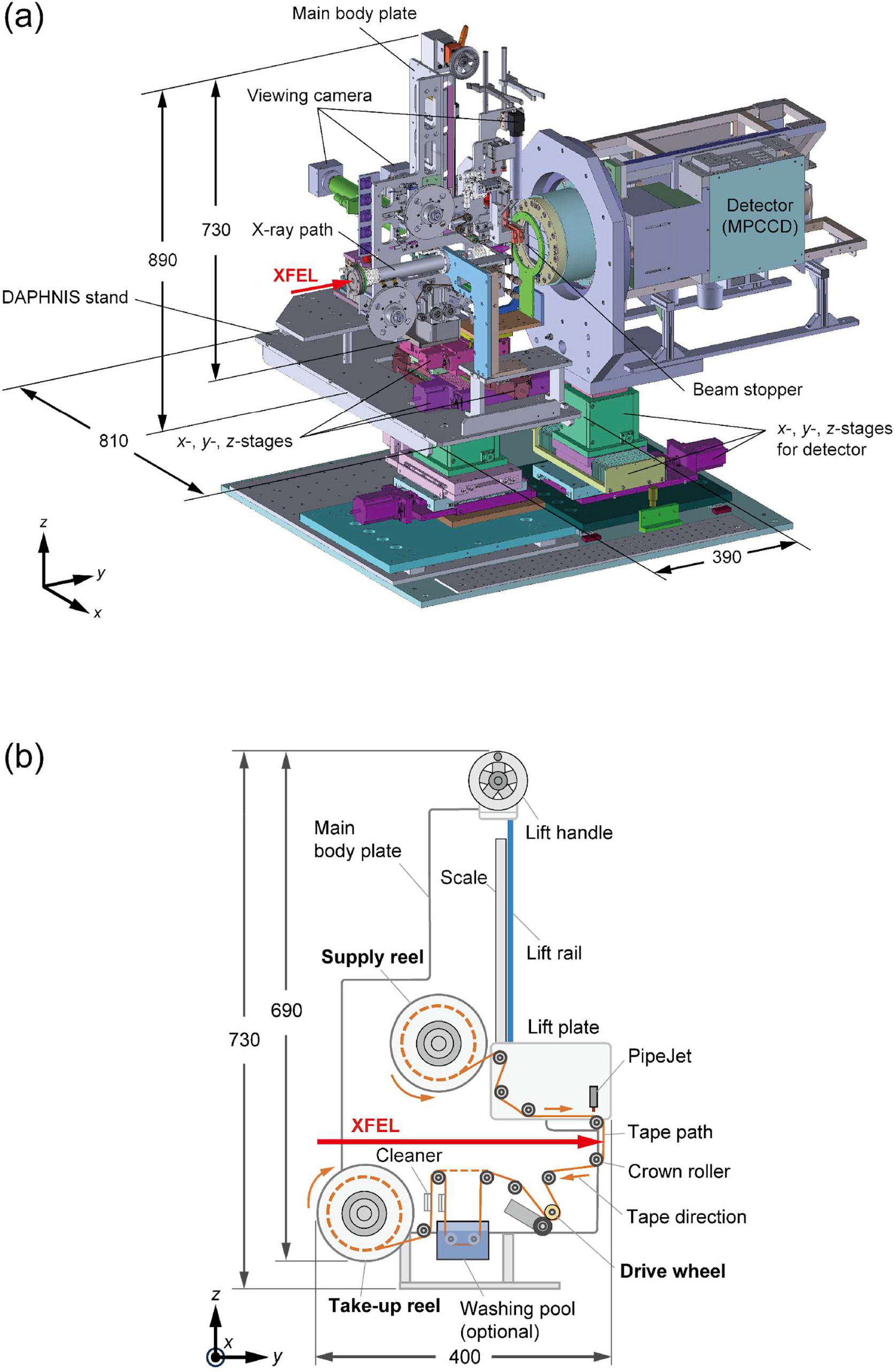
(a) CAD diagram of CoT installed on DAPHNIS. (b) Schematic diagram of the main body plate. Units are in millimeters.

Figure 1b shows a schematic diagram of CoT. A washing pool can be introduced if needed. The reel-to-reel tape drive, which is widely used in serial crystallography at synchrotron facilities (Beyerlein *et al*., 2017, Zielinski *et al*., 2022, Henkel *et al*., 2023), comprises a 100-m-long roll of polyimide film (Kapton®) tape without any water-repellent treatment on the surface, with a thickness of 12.5 µm and width of 3 mm that is inserted into a reel cartridge of two 130-mm-diameter disks (Fig. S1a). The reel cartridge is set on the supply reel unit and rotates counterclockwise to release the tape. The drive wheel comprises a stainless-steel wheel with a diameter of 20 mm and width of 20 mm that is coated by a urethane rubber layer with a thickness of 3 mm, and it is placed at the bottom and rotates clockwise to convey the tape at speeds of 15–300 mm s^−1^. It is possible to set the tape speed to less than 15 mm s^−1^. However, droplets dispensed onto the tape without any water-repellent treatment may not separate. Here, the pulse motor connected to the supply reel applies a weak force in a clockwise direction, which is opposite to the direction of the supply reel’s rotation, to tension the tape. A take-up reel unit with an empty reel cartridge also rotates clockwise to collect the tape at the end of the tape path. Ten crown rollers with a diameter of 16 mm, a width of 7.4 mm, and a curvature radius of 86 mm are installed along the tape path (Fig. S1b). While the tape is fed at a speed of several tens to hundreds of mm s^−1^, these rollers maintain the tape path and suppress the meandering motion of the tape in the *x*-direction in Fig. 1b to within ±0.1 mm.

The commercial piezoelectric injector PipeJet® (Hamilton Company) is used to dispense sample droplets onto the tape surface. Figure 2a shows the droplet ejection mechanism of PipeJet (Streule *et al*., 2004, Hamilton©, 2025). A thin polymer tube with an inner diameter of 200 µm is used as a disposable nozzle that connects the back end to the sample reservoir. A tube with an inner diameter of 200 µm is typically used, but tubes with diameters of 125 µm and 500 µm are also available. A piston connected to a piezostack actuator squeezes the tube over an active area with a length of approximately 5 mm, which displaces the sample solution toward both ends of the tube. The sample solution that is pushed toward the tip is dispensed as nanoliter-scale droplets that form a queue on the tape surface with a constant spacing depending on the tape speed (e.g., a spacing of 1 mm corresponds to a tape speed of 30 mm s^−1^). The volume of the sample droplets depends on the displacement and velocity of the piston, inner diameter of the nozzle, and viscosity of the samples. For pure water, 125, 200, and 500 µm ID tubes can generate between 2–12 nL, 5–18 nL, and 20–75 nL droplets at up to 50 Hz maximum frequency, respectively (Hamilton©, 2025). Figure S2a shows a pure water droplet queue with a volume of 9 nL ejected by the PipeJet with a 200 µm ID tube on the tape surface of the Kapton® film without any water-repellent treatment. Each droplet forms a lowered dome shape with a diameter of about 500 µm and a height of about 90 µm. A commercial 1 mL Terumo syringe is used as the sample reservoir, and the nozzle protrudes 2 mm from the rest of the piezoelectric injector (Fig. S2b). A simple stirring propeller is mounted to the sample reservoir to prevent sedimentation of the microcrystals (Fig. S2c). In the ejection section (Fig. S2d), the nozzle is aimed at the tape surface, and an antistatic brush helps reduce tape charging.

**Figure 2.**
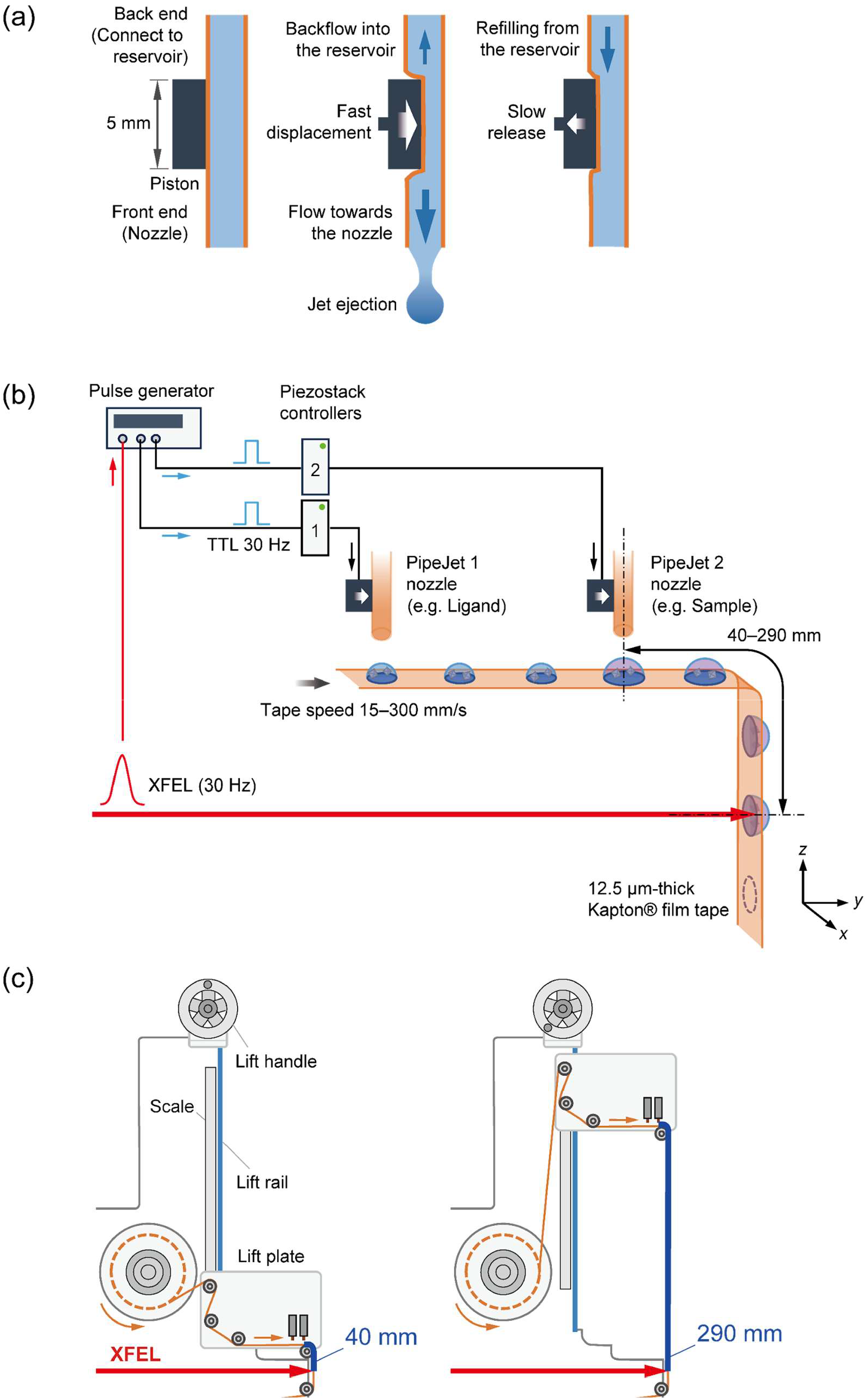
Schematic diagrams of the (a) droplet ejection mechanism, (b) sample delivery synchronized with XFEL irradiation, and (c) control of the distance between the droplet ejection area and XFEL intersecting area.

For mixing experiments, two PipeJet units are mounted to a lift plate: one unit dispenses droplets containing microcrystals, while the other ejects a solution containing a substrate or ligand. The distance between the nozzle tips of PipeJet units 1 and 2 is manually adjusted with respect to the tape speed so that the droplets would overlap precisely, usually by moving PipeJet unit 1 along the *y*-axis direction (e.g., a distance of 15 mm at a tape speed of 30 mm s^−1^). PipeJet units 1 and 2 are individually mounted onto the lift plate with *x-, y*-, and *z*-axis micrometer stages and *x*- and *z*-axis micrometer stages, respectively (Fig. S3). Each droplet is dispensed in synchronization with a transistor-transistor logic (TTL) signal generated by a pulse generator as a trigger signal referenced to the XFEL clock signal at SACLA. The two droplets from the PipeJet units are overlaid, and then irradiated with an XFEL pulse after a certain delay time. As shown in Fig. 2c, the distance from the location where the two droplets are mixed to the XFEL intersecting area can be varied from 40 mm to 290 mm. The lift plate can be moved up and down along the lift rail in the range of 250 mm by turning the lift handle. The delay time after mixing the two droplets can be adjusted from a minimum of 0.13 s to a maximum of 19.3 s depending on the tape speed and distance.

The XFEL pulses are incident perpendicular to the tape broad surface and pass through the tape before reaching the droplets. Positioning the XFEL perpendicular to the tape surface facilitates hitting the crystals in the droplets because it negates the effects of the droplet height and crystal sedimentation. Additionally, the hit ratio of the XFEL pulses onto the droplets is approximately 100% at a speed of 30 mm s^−1^ because the bottom surface of the droplets is wide, allowing the XFEL pulse and the droplets to intersect easily. However, this approach can introduce background noise depending on the thickness of the tape. Figure S4 and S5 show a typical diffraction pattern of HEWL crystals using a 12.5-µm-thick Kapton® tape. Diffraction rings from the tape appear at a low-angle region near the center (5–20 Å resolution). However, the intensity of the rings is sufficiently low (∼10^−2^) compared to those of the diffraction spots derived from the HEWL crystals that they would not influence data processing for structural analysis. Solvent scattering from the droplets is also observed, but the influence of background noise is not significant because the height of the droplet generating the scattering is relatively low (Fig. S4b). The droplets on the tape surface tend to spread since the tape is not treated to water-repellent (Fig. S2a). Assuming the droplets maintain the same contact angle relative to the tape surface, a droplet volume of 10 nL causes a height of approximately 90 µm, while a volume of 14 nL results in approximately 100 µm, resulting in a difference of only approximately 10%.

## 3. Time-resolved mixing SFX experiment

### 3.1. Sample preparation

Mixing experiments were conducted using HEWL as the protein crystals and GlcNAc as an inhibitor. HEWL is a glycoside hydrolase that is widely used in X-ray crystallography as a commercially available model protein and was crystallized by a batch method according to the protocol with modification (Nango *et al*., 2015, Sugahara *et al*., 2015). First, 10 mL of 20 mg mL^−1^ HEWL (FUJIFILM Wako Pure Chemical) dissolved in 0.1 M sodium acetate (pH 3.0) was added to an equivalent volume of crystallization buffer [8% (w/v) PEG 6,000, 4.8 M NaCl, and 0.1 M sodium acetate (pH 3.0)]. The mixture was then equilibrated by using ThermoMixer C (Eppendorf) for 10 min at a rotational speed of 500 rpm and a temperature of 12°C or 17°C, which resulted in HEWL microcrystals with dimensions of 1 or 3–5 µm, respectively. The 1-µm and 3–5-µm crystals had crystal densities of approximately 1.1 × 10^10^ crystals mL^−1^ and 4.8 × 10^8^ crystals mL^−1^, respectively. The microcrystals were then harvested by centrifugation at 3,000 × *g* for 5 min at 4°C. The supernatant was replaced with a harvest buffer (1 M sodium acetate (pH 3.0), 1.7 M NaCl), and the microcrystals were stored at 4°C until use. The 1-µm crystals were diluted at a 1:10 ratio to a final density of ∼1.1 × 10^9^ crystals mL^−1^ using the harvest buffer before SFX measurements while the 3–5-µm crystals were used as is. These densities correspond to 11,000 crystals per drop (10 nL) for the 1-μm crystal and 4,800 crystals per drop for the 3–5-μm crystal. Using a 1.5-μm XFEL beam, it is expected to result in diffraction patterns from a single crystal per image. The protein concentration for each crystal suspension was estimated based on the number of protein molecules in the unit cell and the volume of the crystal.

For the inhibitor, 50 mg mL^−1^ (226 mM) and 100 mg mL^−1^ (452 mM) of GlcNAc were dissolved in water or various buffers including polyethylene glycol (PEG). Then, HEWL crystal droplets were merged with the different inhibitor droplets at different concentrations and buffer types, and a microscope was used to visually check whether the crystal and inhibitor droplets mixed properly. The inhibitor dissolved in water did not seem to mix well with the crystal droplets. Thus, the inhibitor dissolved in a buffer of 1 M NaCl and 15% PEG 4,000 was selected for the mixing experiments. The concentration of the 226 mM GlcNAc solution was approximately 100 and 1,000 times the molar excess relative to the undiluted microcrystal suspension and 10-fold diluted suspension, respectively. The inhibitor solution was filtered through a 0.22-µm filter before measurements.

### 3.2. Diffraction data collection

Data were collected at Experimental Hutch 3 of SACLA beamline 2 (Ishikawa *et al*., 2012, Tono *et al*., 2019) using XFEL pulses with a photon energy of 10 keV, pulse duration of 10 fs or less, and pulse energy of 370 µJ on average before the first mirror in Optics Hutch 1, which contained approximately 2 × 10^11^ photons per pulse at a repetition rate of 30 Hz. The X-ray beam was focused to a full width at half maximum diameter of 1.5 µm by two X-ray ellipsoidal mirrors in the Kirkpatrick– Baez geometry. To avoid the parasitic scattering of X-rays from air, a collimator was mounted on CoT, and helium gas was introduced at a flow rate of 1.4 mL min^−1^. Measurements were performed at a temperature of 25–26°C and relative humidity of 30–40%. Diffraction patterns were recorded by using an MPCCD detector (Phase-III) (Kameshima *et al*., 2014) with eight sensor modules at a sample– detector distance of 70 mm.

Droplets were dispensed at a piezostack stroke of 5% and stroke velocity of 120 µm ms^−1^. A nozzle with an inner diameter of 200 µm was mounted to each PipeJet unit. One PipeJet unit ejected an inhibitor droplet containing GlcNAc, which was overlaid by an equal volume droplet containing HEWL crystals ejected by the second PipeJet unit. Individually, the crystal and inhibitor droplets had an average volume of 10–14 nL with a diameter of about 520–580 µm and a height of about 90–100 µm on the tape surface. The merged droplet reached a diameter of about 660–740 µm and a height of about 115– 130 µm. The tape speed was set to 30 mm s^−1^ while the distance between the point at which the second droplet is ejected and XFEL irradiation point was varied in the range of 40–290 mm. Thus, the mixing time between the merging of the two droplets and XFEL irradiation was varied in the range of 1.3–9.7s. Finally, 13 datasets were obtained under different conditions including the resting states of the HEWL crystals without the addition of inhibitor droplets. These data were collected during 36 hours of beamtime under proposal number 2023A8008.

### 3.3. Data processing and structure determination

Diffraction images were filtered to extract “hit images” that included diffraction patterns from the crystals by using Cheetah (Barty *et al*., 2014), which was modified for processing SFX data at SACLA (Nakane *et al*., 2016). A hit image was defined as containing a minimum of 20 Bragg peaks. The detector geometry was refined by geoptimizer in CrystFEL (version 0.10.2) (White *et al*., 2013). In addition, autoindexing and integration were performed by using CrystFEL. The resting structure of lysozyme was solved by using the molecular replacement method in the software Phaser (McCoy *et al*., 2007). The PDB model 6jzi (Shimazu *et al*., 2019) from the PDB database was applied as a reference after the water molecules and metal ions were removed. Further refinement of the structure was conducted using Phenix (Adams *et al*., 2010) and COOT (Emsley *et al*., 2010). Finally, MOLPROBITY (Chen *et al*., 2010) was used for data validation. The differences in electron density maps between the mixing data and resting state data were calculated by the CCP4 toolkit (Agirre *et al*., 2023). All structure images were created by using Pymol (Schrödinger & DeLano, 2020).

## 4. Results

To demonstrate the performance of the compact tape drive (CoT), we conducted time-resolved SFX experiments mixing HEWL crystals of different sizes and the inhibitor GlcNAc. Two concentrations of the inhibitor were prepared and first dispensed as a droplet of 10–14 nL on the tape. Subsequently, HEWL crystals were ejected into the first droplet in the same volume, which was basically mixed by diffusion. The equilibration/reaction time was adjusted by moving the lift plate on CoT. The 1-µm HEWL crystals were mixed with 226 mM GlcNAc and measured at delay times of 1.3, 5.0, 7.5, and 9.7 s or mixed with 452 mM GlcNAc and measured at delay times of 1.3, 2.5, 4.0, and 5.0 s. The hit ratio was 14–31%, and the indexed rate was 77% on average. The 3–5-µm HEWL crystals were mixed with 226 mM GlcNAc and measured at delay times of 2.0, 5.0, and 9.7 s. The hit ratio was 40–54%, and the indexed rate was 80% on average. Each dataset comprised 10,000–24,000 indexed images. Refined structural models were determined with a resolution of 1.68–1.82 Å. One dataset typically required approximately one hour to collect and used one roll of tape spooling out at 30 mm s^−1^. Data collection proceeded smoothly and was often only interrupted if/when tape replacement was required. This setup is basically intended to be operated by the user. Thus, the users need to replace a pair of reel-to-reel tapes. Tape replacement typically takes about 5 minutes, although it may take longer initially. In the experiments, 10–14 nL of the crystal suspension was used per droplet. Thus, the sample consumption for the 1-µm HEWL crystals was 0.1 µg per diffraction image, or 0.5–1.0 mg for 10,000 indexed images. The sample consumption for the 3–5-µm HEWL crystals was 2 µg per diffraction image, or 12–17 mg for 10,000 indexed images.

The resting state structures of the 1-µm and 3–5-µm HEWL crystals both show an acetate ion and several water molecules in the active site (Table 1). To visualize the binding process of the inhibitor, difference electron density (DED) maps were calculated by subtracting the resting state data from the time-resolved mixing SFX data. The *F*_o_ – *F*_c_ maps show not only the electron densities corresponding to the inhibitor but also the original acetate ion and water molecules, which prevents an accurate estimation of the inhibitor binding ratio. Therefore, the inhibitor binding ratio was evaluated based on the intensity of the peaks in the DED maps. Figure 3 shows the DED maps of 3–5 µm HEWL crystals mixed with 226 mM GlcNAc. At a mixing time of 2.0 s, the inhibitor was partially bound to the active site. The inhibitor binding ratio increased at a mixing time of 5.0 s but then remained stable as the mixing time increased further to 9.7 s, as presented in Table 2. Figure 4 shows the DED maps of 1-µm HEWL crystals mixed with 226 mM GlcNAc, which indicates that the inhibitor was partially bound to the active site at the mixing time of 1.3 s. Despite the shorter mixing time compared to the 3–5 µm HEWL crystals, the inhibitor binding ratio was the same, as given in Table 3. In addition, the inhibitor binding ratio gradually increased over time. Figure 5 shows the DED maps of 1-µm HEWL crystals mixed with 452 mM GlcNAc. At a mixing time of 1.3 s, the peak intensity at the active site is higher than that shown in Fig. 4, which suggests that increasing the inhibitor concentration increased the inhibitor binding ratio. As given in Table 4, the inhibitor binding ratio also gradually increased with the mixing time. In summary, we successfully observed an increase in the inhibitor binding ratio according to the crystal size and mixing time.

**Table 1.**
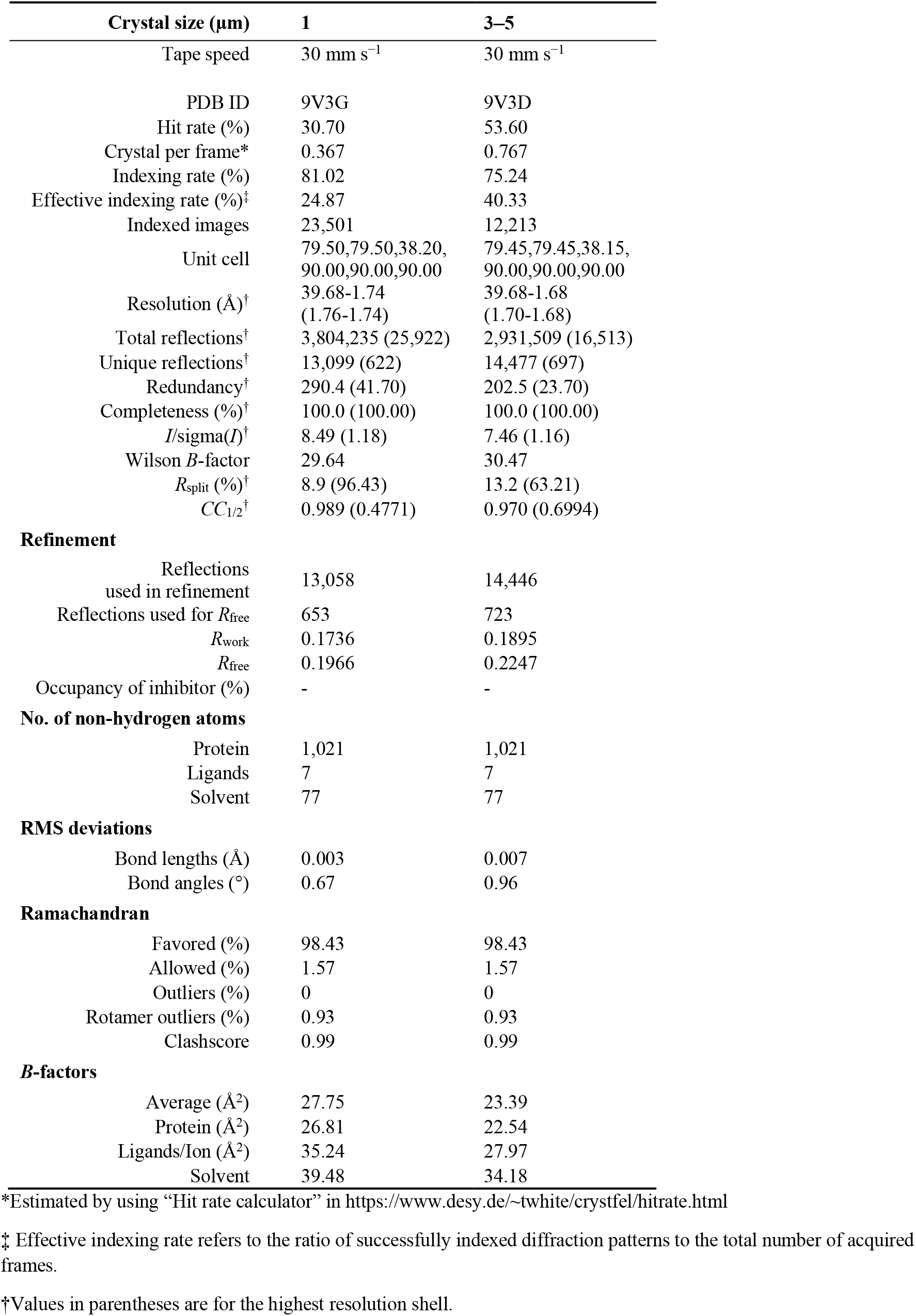
Crystallographic statistics of 1-µm and 3–5-µm HEWL crystals without the inhibitor.

**Table 2.**
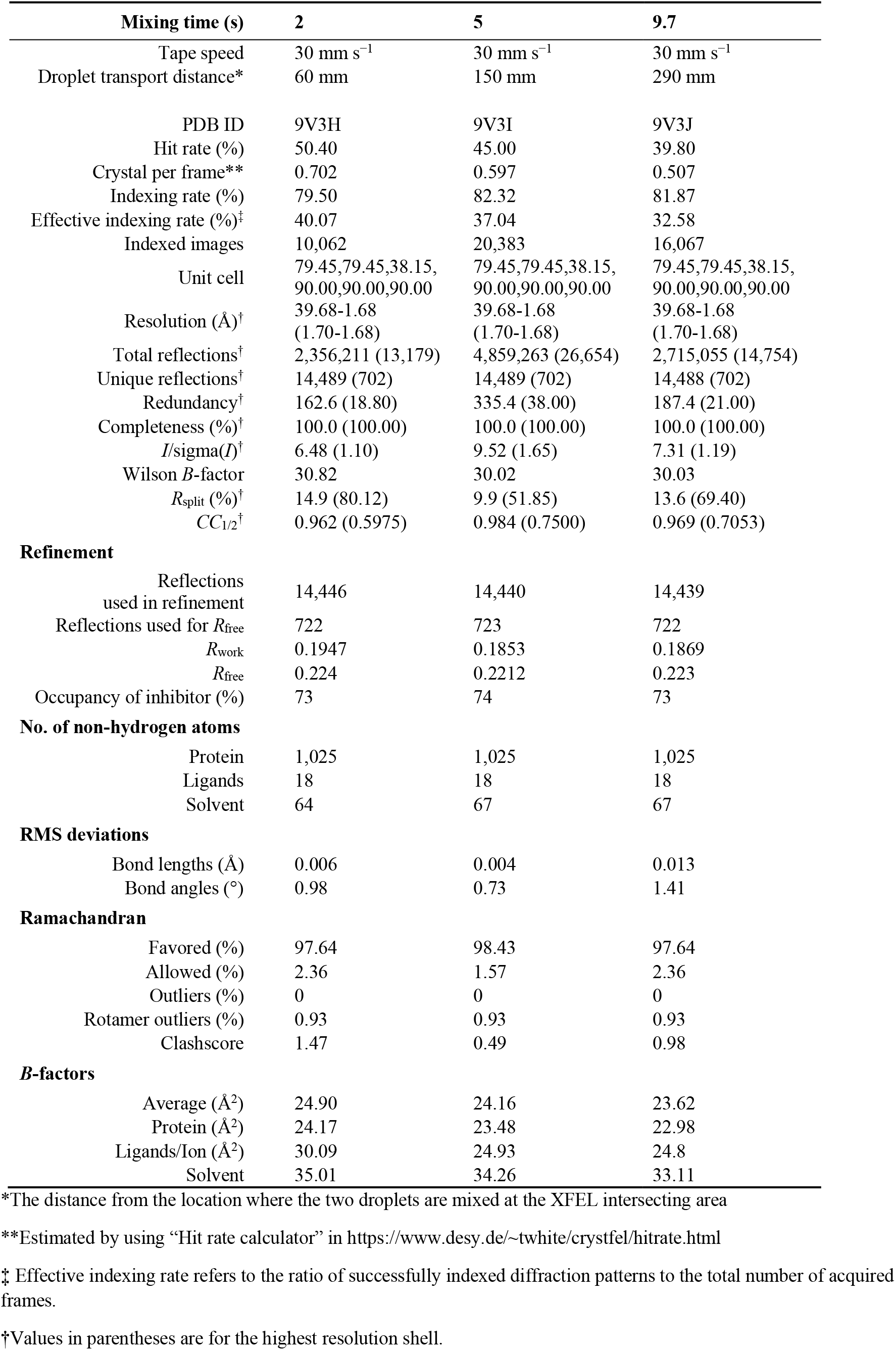
Crystallographic statistics of 3–5-µm HEWL crystals bound with 226 mM GlcNAc.

**Table 3.** Crystallographic statistics of 1-µm HEWL crystals bound with 226 mM GlcNAc.

| Mixing time (s) | 1.3 | 5 | 7.5 | 9.7 |
| --- | --- | --- | --- | --- |
| Tape speed | 30 mm s<sup>-1</sup> | 30 mm s<sup>-1</sup> | 30 mm s<sup>-1</sup> | 30 mm s<sup>-1</sup> |
| Droplet transport distance* | 40 mm | 150 mm | 225 mm | 290 mm |
| PDB ID | 9V3K | 9V3L | 9V3M | 9V3N |
| Hit rate (%) | 25.77 | 23.50 | 24.20 | 14.40 |
| Crystal per frame** | 0.298 | 0.268 | 0.277 | 0.156 |
| Indexing rate (%) | 80.84 | 64.07 | 81.98 | 80.19 |
| Effective indexing rate (%)‡ | 20.83 | 15.06 | 19.84 | 11.55 |
| Indexed images | 39,020 | 16,967 | 11,967 | 17407 |
| Unit cell | 79.50,79.50,38.20,<br>90.00,90.00,90.00 | 79.50,79.50,38.20,<br>90.00,90.00,90.00 | 79.50,79.50,38.20,<br>90.00,90.00,90.00 | 79.50,79.50,38.20,<br>90.00,90.00,90.00 |
| Resolution (Å)† | 39.68-1.72<br>(1.74-1.72) | 39.68-1.75<br>(1.78-1.75) | 39.68-1.77<br>(1.80-1.77) | 39.68-1.78<br>(1.81-1.78) |
| Total reflections† | 6,390,960 (33,815) | 2,927,612 (19,507) | 2,152,845 (19,216) | 2,882,272 (18,231) |
| Unique reflections† | 13,560 (661) | 12,892 (611) | 12,477 (620) | 12,281 (613) |
| Redundancy† | 471.3 (51.20) | 227.1 (31.90) | 172.5 (31.00) | 234.7 (29.70) |
| Completeness (%)† | 100.0 (100.00) | 100.0 (100.00) | 100.0 (100.00) | 100.0 (100.00) |
| *I*/$\sigma$(*I*)† | 10.63 (1.03) | 7.56 (1.02) | 6.85 (1.04) | 8.43 (1.32) |
| Wilson *B*-factor | 29.81 | 28.58 | 29.23 | 28.71 |
| *R*<sub>split</sub> (%)† | 6.9 (101.02) | 10.8 (105.96) | 11.5 (99.77) | 9.1 (85.43) |
| *CC*<sub>1/2</sub>† | 0.994 (0.4427) | 0.985 (0.4146) | 0.980 (0.4291) | 0.989 (0.4833) |
| **Refinement** |  |  |  |  |
| Reflections used in refinement | 13,518 | 12,850 | 12,437 | 12,239 |
| Reflections used for *R*<sub>free</sub> | 677 | 643 | 622 | 612 |
| *R*<sub>work</sub> | 0.1774 | 0.1788 | 0.1821 | 0.1672 |
| *R*<sub>free</sub> | 0.2056 | 0.1988 | 0.2185 | 0.1882 |
| Occupancy of inhibitor (%) | 68 | 74 | 77 | 79 |
| **No. of non-hydrogen atoms** |  |  |  |  |
| Protein | 1,025 | 1,025 | 1,025 | 1,015 |
| Ligands | 18 | 18 | 18 | 18 |
| Solvent | 64 | 64 | 64 | 67 |
| **RMS deviations** |  |  |  |  |
| Bond lengths (Å) | 0.003 | 0.005 | 0.003 | 0.007 |
| Bond angles (°) | 0.67 | 0.84 | 0.67 | 0.89 |
| **Ramachandran** |  |  |  |  |
| Favored (%) | 98.43 | 99.21 | 98.43 | 99.21 |
| Allowed (%) | 1.57 | 0.79 | 1.57 | 0.79 |
| Outliers (%) | 0 | 0 | 0 | 0 |
| Rotamer outliers (%) | 1.87 | 0.93 | 0.93 | 0.95 |
| Clashscore | 0.98 | 1.47 | 0.49 | 0.99 |
| ***B*-factors** |  |  |  |  |
| Average (Å<sup>2</sup>) | 28.02 | 27.19 | 26.61 | 27.35 |
| Protein (Å<sup>2</sup>) | 27.27 | 26.65 | 26.10 | 26.73 |
| Ligands/Ion (Å<sup>2</sup>) | 37.63 | 29.67 | 28.96 | 28.27 |
| Solvent | 37.34 | 35.14 | 34.22 | 36.56 |

\*The distance from the location where the two droplets are mixed at the XFEL intersecting area
\*\*Estimated by using “Hit rate calculator” in https://www.desy.de/~twhite/crystfel/hirate.html
‡ Effective indexing rate refers to the ratio of successfully indexed diffraction patterns to the total number of acquired frames.
†Values in parentheses are for the highest resolution shell.

**Table 4.** Crystallographic statistics of 1-µm HEWL crystals bound with 452 mM GlcNAc.

| Mixing time (s) | 1.3 | 2.5 | 4 | 5 |
| --- | --- | --- | --- | --- |
| Tape speed | 30 mm s<sup>-1</sup> | 30 mm s<sup>-1</sup> | 30 mm s<sup>-1</sup> | 30 mm s<sup>-1</sup> |
| Droplet transport distance* | 40 mm | 75 mm | 120 mm | 150 mm |
| PDB ID | 9V3O | 9V3P | 9V3Q | 9V3R |
| Hit rate (%) | 25.70 | 24.88 | 21.50 | 15.20 |
| Crystal per frame** | 0.297 | 0.286 | 0.242 | 0.165 |
| Indexing rate (%) | 82.36 | 82.45 | 73.96 | 78.74 |
| Effective indexing rate (%)† | 21.17 | 20.51 | 15.90 | 11.97 |
| Indexed images | 15,331 | 12,438 | 12,723 | 17,626 |
| Unit cell | 79.50,79.50,38.20,<br>90.00,90.00,90.00 | 79.50,79.50,38.20,<br>90.00,90.00,90.00 | 79.50,79.50,38.20,<br>90.00,90.00,90.00 | 79.50,79.50,38.20,<br>90.00,90.00,90.00 |
| Resolution (Å)† | 39.68-1.80<br>(1.83-1.80) | 39.68-1.81<br>(1.84-1.81) | 39.68-1.80<br>(1.83-1.80) | 39.68-1.82<br>(1.85-1.82) |
| Total reflections† | 2,451,958 (20,282) | 1,782,793 (16,506) | 1,901,023 (14,302) | 1,689,033 (15,393) |
| Unique reflections† | 11,866 (570) | 11,685 (572) | 11,866 (570) | 11,492 (552) |
| Redundancy† | 206.6 (35.60) | 152.6 (28.90) | 160.2 (25.10) | 147.0 (27.90) |
| Completeness (%)† | 100.0 (100.00) | 100.0 (100.00) | 100.0 (100.00) | 100.0 (100.00) |
| *I*/$\sigma$(*I*)† | 7.46 (1.15) | 6.66 (1.18) | 6.80 (1.13) | 6.62 (1.12) |
| Wilson *B*-factor | 29.49 | 28.82 | 29.02 | 28.59 |
| *R*<sub>split</sub> (%)† | 10.2 (91.35) | 12.1 (94.09) | 11.4 (98.65) | 12.1 (96.06) |
| *CC*<sub>1/2</sub>† | 0.985 (0.4231) | 0.979 (0.4079) | 0.980 (0.4280) | 0.979 (0.4353) |
| **Refinement** |  |  |  |  |
| Reflections used in refinement | 11,825 | 11,645 | 11,823 | 11,451 |
| Reflections used for *R*<sub>free</sub> | 592 | 583 | 592 | 573 |
| *R*<sub>work</sub> | 0.1712 | 0.1728 | 0.1773 | 0.1727 |
| *R*<sub>free</sub> | 0.2082 | 0.2218 | 0.2161 | 0.1947 |
| Occupancy of inhibitor (%) | 75 | 77 | 81 | 79 |
| **No. of non-hydrogen atoms** |  |  |  |  |
| Protein | 1,025 | 1,025 | 1,025 | 1,025 |
| Ligands | 18 | 18 | 18 | 18 |
| Solvent | 64 | 64 | 64 | 64 |
| **RMS deviations** |  |  |  |  |
| Bond lengths (Å) | 0.003 | 0.006 | 0.003 | 0.006 |
| Bond angles (°) | 0.71 | 0.92 | 0.66 | 0.86 |
| **Ramachandran** |  |  |  |  |
| Favored (%) | 99.21 | 99.21 | 99.21 | 98.43 |
| Allowed (%) | 0.79 | 0.79 | 0.79 | 1.57 |
| Outliers (%) | 0 | 0 | 0 | 0 |
| Rotamer outliers (%) | 0.93 | 0.93 | 0.93 | 0.93 |
| Clashscore | 1.47 | 1.47 | 0.98 | 0.98 |
| ***B*-factors** |  |  |  |  |
| Average (Å<sup>2</sup>) | 28.83 | 28.00 | 26.86 | 27.14 |
| Protein (Å<sup>2</sup>) | 28.17 | 27.36 | 26.39 | 26.72 |
| Ligands/Ion (Å<sup>2</sup>) | 35.72 | 33.26 | 26.95 | 26.6 |
| Solvent | 37.49 | 36.76 | 34.21 | 33.97 |

\*The distance from the location where the two droplets are mixed at the XFEL intersecting area
\*\*Estimated by using “Hit rate calculator” in https://www.desy.de/~twhite/crystfel/hirate.html
† Effective indexing rate refers to the ratio of successfully indexed diffraction patterns to the total number of acquired frames.
† Values in parentheses are for the highest resolution shell.

**Figure 3.**
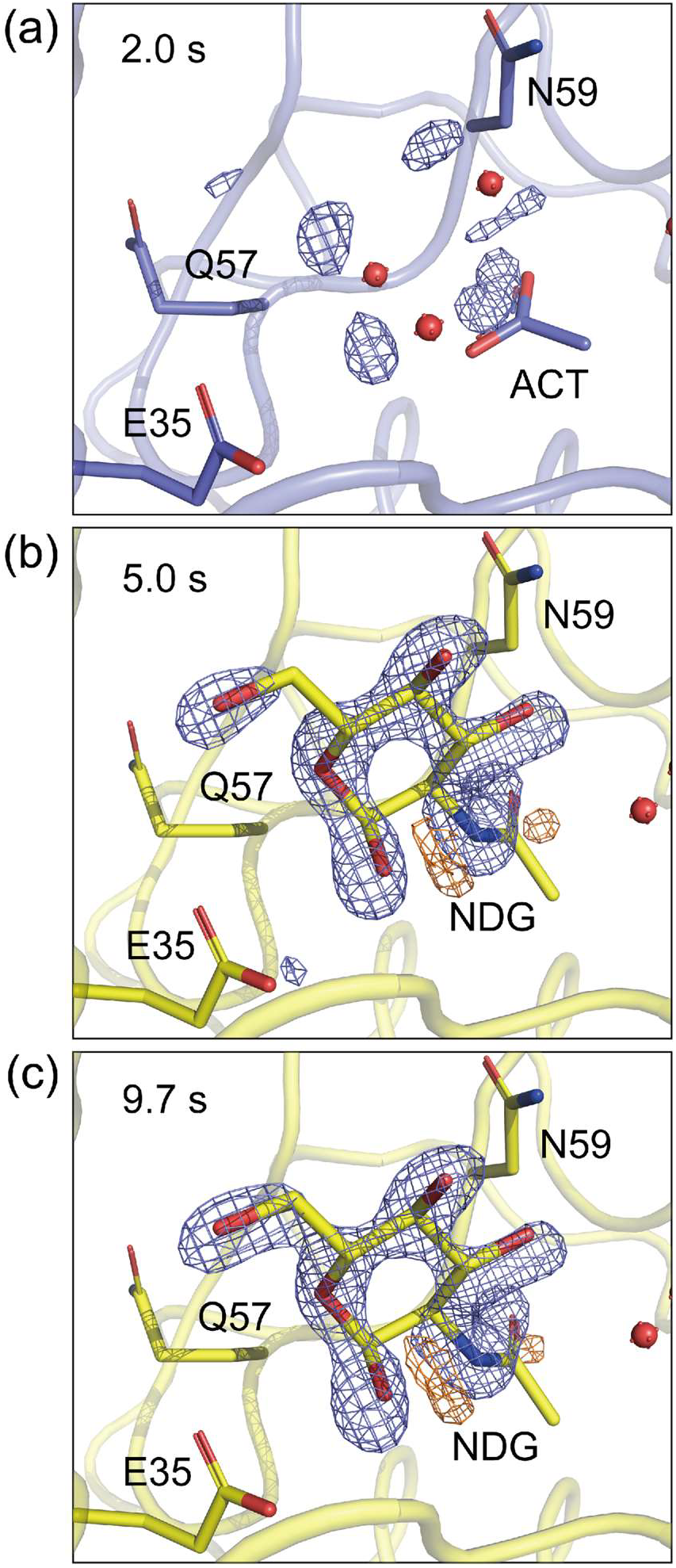
*F*_o (bound)_ – *F*_o (resting)_ difference electron density maps obtained by mixing 3–5-µm HEWL crystals with 226 mM GlcNAc at mixing times of (a) 2.0 s, (b) 5.0 s, and (c) 9.7 s. The maps are contoured at 5.00 sigma. The HEWL structure in the absence of GlcNAc is shown with blue carbon atoms while the GlcNAc-bound HEWL structure is shown with yellow carbon atoms. The positive and negative electron density peaks are depicted in slate and TV-yellow, respectively.

**Figure 4.**
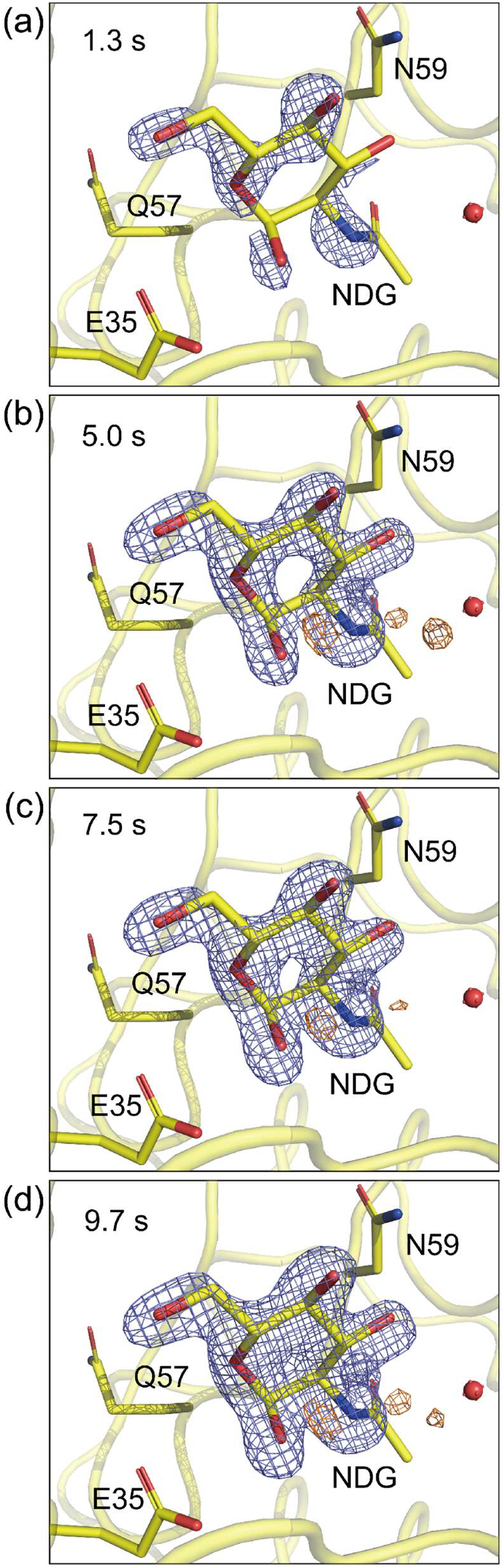
*F*_o (bound)_ – *F*_o (resting)_ difference electron density maps obtained by mixing 1-µm HEWL crystals with 226 mM GlcNAc at mixing times of (a) 1.3 s, (b) 5.0 s, (c) 7.5 s, and (d) 9.7 s. The maps are contoured at 6.00 sigma. The GlcNAc-bound HEWL structure is shown with yellow carbon atoms. The positive and negative electron density peaks are depicted in slate and TV-yellow, respectively.

**Figure 5.**
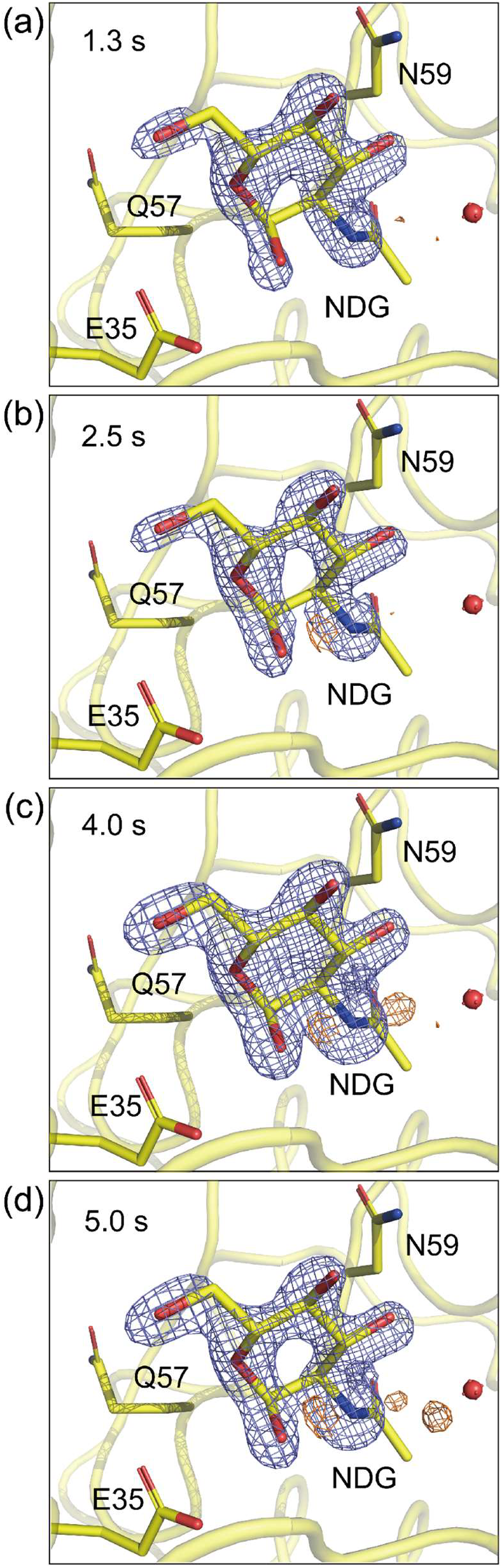
*F*_o (bound)_ – *F*_o (resting)_ difference electron density maps obtained by mixing 1-µm HEWL crystals with 452 mM GlcNAc at mixing times of (a) 1.3 s, (b) 2.5 s, (c) 4.0 s, and (d) 5.0 s. The maps are contoured at 6.00 sigma. The GlcNAc-bound HEWL structure is shown with yellow carbon atoms. The positive and negative electron density peaks are depicted in slate and TV-yellow, respectively.

## 5. Discussion

The main differences between CoT and Fuller *et al*.’s DOT method are that the XFEL pulses pass through the tape and that piezoelectric injectors are used rather than acoustic droplets to dispense droplets. In the experiments performed at SACLA in 2023, the volume of droplets was three to ten times larger than those typically obtained with the DOT method, but the sample volume can be reduced. After conducting tests, we found that the volume depends on the condition of the device and nozzle. Subsequently, we successfully ejected droplets of 4 to 6 nL using a 200 μm nozzle with a solution containing crystals. With a 125 μm nozzle, we were able to dispense droplets including buffer or crystals of 1–2 nL. One concern is that the tape may be punctured by the intense XFEL pulses, which matters mostly for systems that clean and reuse the same tape and less for reel-to-reel systems. In fact, we observed small holes in the tape surface after XFEL irradiation that are fully consistent with the X-ray beam size and alignment setup (Fig. S6). However, the tape was driven stably without any problems during data collection indicating that small perforations drilled through by the 1.5 µm XFEL beam diameter did not compromise the CoT. Although the tape was replaced after a single-use, it can potentially be reused by shifting the XFEL intersecting area and installing a cleaning bath to remove salts and crystals. The background noise induced by the tape was relatively low because the tape was only 12.5 µm thick, which is much thinner than the sample streams from the high-viscosity sample injector typically used at SACLA, which are 75–100 µm in diameter (Fig. S4).

In the experiments, the crystal and inhibitor droplets had the same volumes, so mixing them halved the inhibitor concentrations from 226 mM to 113 mM and from 452 mM to 226 mM. Given that GlcNAc has an affinity constant of 47.6 mM (Kumagai *et al*., 1992), these concentrations are sufficient. Inhibitor binding was observed after a mixing time of 1.3 s under all conditions, and the bound inhibitors showed different occupancies depending on the concentration and crystal size. According to our knowledge, this is the first report to perform time-resolved mixing SFX experiments using different HEWL crystal sizes and inhibitor concentrations. However, to fully characterize the time-resolved crystallography parameter space for enzyme reactions, a comprehensive study should include i) a range of homogenous crystal-size slurries (e.g. 0.5, 1, 2, 3, 5, 7.5, and 10 micron dimensions), ii) a range of ligand concentrations (e.g. from 0.1× to 10×) related to the known *K*_M_, *K*_I_, or *K*_D_ values measured in solutions and correlated to crystals, iii) known or experimentally determined ligand diffusion times in water and mother liquor, as well as through crystal lattices, iv) across a range of temperatures, and v) more than one space group and crystal packing conditions across the same reaction coordinate. The diffusion time of small molecules into crystals is known to depend on the crystal size and ligand concentration, which we successfully demonstrated experimentally using CoT. CoT is capable of observing mixing times of 0.1–19.3 s, and it is possible that binding could have taken place at an earlier mixing time. However, we did not test such conditions because of the limited beamtime available at SACLA. Butryn *et al*. collected time-resolved SFX data at a mixing and equilibration time of 0.6 s using 3–5-µm HEWL crystals with 16.7 mM GlcNAc as the final concentration (Butryn *et al*., 2021). They injected multiple, picoliter volume droplets of the inhibitor solution into a ∼3 nL crystal slurry droplet to create turbulence, which resulted in a higher mixing efficiency. In this study, we induced rapid diffusion by increasing the inhibitor concentration by 3–14 times compared with that used by Butryn *et al*. (43.7 mM) to achieve inhibitor binding at a similar mixing time. If increasing the ligand concentration is difficult because of constraints such as its solubility, alternative approaches to accelerate ligand diffusion into crystals would be necessary such as turbulent mixing or heating. The volume of droplets also affects the diffusion time. Therefore, reducing the volume is effective for rapid diffusion, and the use of a small nozzle such as 125 µm are recommended.

In time-resolved SFX, sample consumption is a critical concern because protein and ligand samples are often scarce and/or expensive. Moreover, for a TR-SFX reaction coordinate, data need to be collected at multiple time points to visualize the structural changes over time. We summarized each sample consumption used in the mixing experiments as shown in Table S1–S3. With CoT, collecting 10,000 indexed images of 3–5-µm HEWL crystals consumed 12–21 mg of HEWL. Meanwhile, Butryn *et al*. reported that they consumed 6.5–27 mg of HEWL to collect 10,000 indexed images of the same crystal size. When we used 1-µm HEWL crystals, we only used 0.4–1 mg of HEWL to obtain 10,000 indexed images, which indicates that the sample consumption can be decreased by reducing the crystal size. Furthermore, we used a droplet volume of 10–14 nL in the experiments, which can also be reduced to improve efficiency. The droplet volume ejected by the PipeJet unit depends on the size of the nozzle and the composition and viscosity of the solution. Offline tests with CoT using a nozzle size of 200 µm showed that it could dispense GlcNAc droplets with a volume of 3–6 nL and HEWL crystal droplets with a volume of 4–6 nL by optimizing the parameters for droplet ejection. A thinner nozzle of 125 µm can be used to further reduce the droplet volume to 1–2 nL.

## 6. Conclusion

In this study, we developed the compact tape-driven sample delivery system, CoT. Piezoelectric injectors were used to dispense sample droplets, and XFEL pulses perpendicularly irradiated to the tape to facilitate alignment of the intersection of the sample droplets with the XFEL. The tape transport speed and the distance between the droplet ejection area and the XFEL intersecting area were designed to be variable, allowing the mixing time range from 0.1 to 19.3 s. We successfully demonstrated that CoT can be employed for time-resolved mixing SFX experiments using HEWL crystals of different crystal sizes and solution concentrations. Under all conditions, the inhibitor was observed to bind within a mixing time of 1.3 s. Reducing the HEWL crystal size to 1 µm was shown to decrease the sample consumption to less than 1 mg for 10,000 indexed images. It is also applicable to light-triggered pump-probe SFX because a pump laser can be introduced to the setup. CoT is expected to reduce the sample consumption of time-resolved SFX experiments and to contribute greatly to the dynamic structural analysis of various proteins in the future.

## Acknowledgements

We acknowledge the members of the Engineering Team of the RIKEN SPring-8 Center for their technical support. We acknowledge the computational support from the SACLA HPC system and computational resources of SACLA HPC provided by RIKEN through the HPCI System Research Project (Project ID: hp230365, hp240364). XFEL experiments were conducted at BL2 of SACLA with the approval of the Japan Synchrotron Radiation Research Institute (JASRI) (Proposal Numbers 2017B8038, 2018B8056, 2018A8042, 2018A8046, 2021A8015, 2022A8026, and 2023A8008). J.K. acknowledges S. Owada for the fruitful discussion.

## Author Contributions

K.T., A.M.O., S.I., E.N., and M.Y. conceived the research; J.K., Y.S., A.Y., T.T., Y.I., K.T., and E.N. developed the instrumentation; A.Y., T.T., N.N., and E.N. prepared the HEWL microcrystals and inhibitor; F.L. processed the data; F.L. and E.N. analyzed the data; J.K. and E.N. considered the operation of the system. J.K., F.L., A.Y., T.T., N.N., and E.N. performed data collection. J.K., F.L., N.N., and E.N. wrote the manuscript.

## Data Availability

Coordinates and structure factors that were generated during the course of this study have been deposited in the Protein Data Bank with the accession codes 9V3D, 9V3G, 9V3H, 9V3I, 9V3J, 9V3K, 9V3L, 9V3M, 9V3N, 9V3O, 9V3P, 9V3Q, and 9V3R. Raw diffraction data has been deposited in the CXIDB entry #236 (doi.org/10.11577/2572337).

## Funding Information

K.T., S.I., and E.N. acknowledge financial support from the X-ray Free-Electron Laser Priority Strategy Program from the Ministry of Education, Culture, Sports, Science and Technology (MEXT). S.I. and E.N. acknowledge the Platform Project for Supporting Drug Discovery and Life Science Research [Basis for Supporting Innovative Drug Discovery and Life Science Research (BINDS)] from the Japan Agency for Medical Research and Development (AMED), and the SACLA/SPring-8 Basic Development Program from the RIKEN SPring-8 Center. E.N. acknowledges financial support from Takeda Science Foundation. A.M.O. acknowledges financial support in part, from a Wellcome Investigator Award (award No. 210734/Z/18/Z), a Royal Society Wolfson Fellowship (award No. RSWF\R2\182017), and a UKRI International Science Partnerships Fund (award No. ISPF-229).

## Notes

https://doi.org/10.11577/2572337

### Competing Interest Statement

The authors have declared no competing interest.

## References

Adams, P. D., Afonine, P. V., Bunkoczi, G., Chen, V. B., Davis, I. W., Echols, N., Headd, J. J., Hung, L. W., Kapral, G. J., Grosse-Kunstleve, R. W., McCoy, A. J., Moriarty, N. W., Oeffner, R., Read, R. J., Richardson, D. C., Richardson, J. S., Terwilliger, T. C. & Zwart, P. H. (2010). Acta Crystallogr D Biol Crystallogr 66, 213–221.

Agirre, J., Atanasova, M., Bagdonas, H., Ballard, C. B., Basle, A., Beilsten-Edmands, J., Borges, R. J., Brown, D. G., Burgos-Marmol, J. J., Berrisford, J. M., Bond, P. S., Caballero, I., Catapano, L., Chojnowski, G., Cook, A. G., Cowtan, K. D., Croll, T. I., Debreczeni, J. E., Devenish, N. E., Dodson, E. J., Drevon, T. R., Emsley, P., Evans, G., Evans, P. R., Fando, M., Foadi, J., Fuentes-Montero, L., Garman, E. F., Gerstel, M., Gildea, R. J., Hatti, K., Hekkelman, M. L., Heuser, P., Hoh, S. W., Hough, M. A., Jenkins, H. T., Jimenez, E., Joosten, R. P., Keegan, R. M., Keep, N., Krissinel, E. B., Kolenko, P., Kovalevskiy, O., Lamzin, V. S., Lawson, D. M., Lebedev, A. A., Leslie, A. G. W., Lohkamp, B., Long, F., Maly, M., McCoy, A. J., McNicholas, S. J., Medina, A., Millan, C., Murray, J. W., Murshudov, G. N., Nicholls, R. A., Noble, M. E. M., Oeffner, R., Pannu, N. S., Parkhurst, J. M., Pearce, N., Pereira, J., Perrakis, A., Powell, H. R., Read, R. J., Rigden, D. J., Rochira, W., Sammito, M., Sanchez Rodriguez, F., Sheldrick, G. M., Shelley, K. L., Simkovic, F., Simpkin, A. J., Skubak, P., Sobolev, E., Steiner, R. A., Stevenson, K., Tews, I., Thomas, J. M. H., Thorn, A., Valls, J. T., Uski, V., Uson, I., Vagin, A., Velankar, S., Vollmar, M., Walden, H., Waterman, D., Wilson, K. S., Winn, M. D., Winter, G., Wojdyr, M. & Yamashita, K. (2023). Acta Crystallogr D Struct Biol 79, 449–461.

Barends, T. R. M., Stauch, B., Cherezov, V. & Schlichting, I. (2022). Nat Rev Methods Primers 2.

Barty, A., Kirian, R. A., Maia, F. R., Hantke, M., Yoon, C. H., White, T. A. & Chapman, H. (2014). J Appl Crystallogr 47, 1118–1131.

Beyerlein, K. R., Dierksmeyer, D., Mariani, V., Kuhn, M., Sarrou, I., Ottaviano, A., Awel, S., Knoska, J., Fuglerud, S., Jonsson, O., Stern, S., Wiedorn, M. O., Yefanov, O., Adriano, L., Bean, R., Burkhardt, A., Fischer, P., Heymann, M., Horke, D. A., Jungnickel, K. E. J., Kovaleva, E., Lorbeer, O., Metz, M., Meyer, J., Morgan, A., Pande, K., Panneerselvam, S., Seuring, C., Tolstikova, A., Lieske, J., Aplin, S., Roessle, M., White, T. A., Chapman, H. N., Meents, A. & Oberthuer, D. (2017). IUCrJ 4, 769–777.

Butryn, A., Simon, P. S., Aller, P., Hinchliffe, P., Massad, R. N., Leen, G., Tooke, C. L., Bogacz, I., Kim, I. S., Bhowmick, A., Brewster, A. S., Devenish, N. E., Brem, J., Kamps, J., Lang, P. A., Rabe, P., Axford, D., Beale, J. H., Davy, B., Ebrahim, A., Orlans, J., Storm, S. L. S., Zhou, T., Owada, S., Tanaka, R., Tono, K., Evans, G., Owen, R. L., Houle, F. A., Sauter, N. K., Schofield, C. J., Spencer, J., Yachandra, V. K., Yano, J., Kern, J. F. & Orville, A. M. (2021). Nat Commun 12, 4461.

Chapman, H. N., Fromme, P., Barty, A., White, T. A., Kirian, R. A., Aquila, A., Hunter, M. S., Schulz, J., DePonte, D. P., Weierstall, U., Doak, R. B., Maia, F. R., Martin, A. V., Schlichting, I., Lomb, L., Coppola, N., Shoeman, R. L., Epp, S. W., Hartmann, R., Rolles, D., Rudenko, A., Foucar, L., Kimmel, N., Weidenspointner, G., Holl, P., Liang, M., Barthelmess, M., Caleman, C., Boutet, S., Bogan, M. J., Krzywinski, J., Bostedt, C., Bajt, S., Gumprecht, L., Rudek, B., Erk, B., Schmidt, C., Homke, A., Reich, C., Pietschner, D., Struder, L., Hauser, G., Gorke, H., Ullrich, J., Herrmann, S., Schaller, G., Schopper, F., Soltau, H., Kuhnel, K. U., Messerschmidt, M., Bozek, J. D., Hau-Riege, S. P., Frank, M., Hampton, C. Y., Sierra, R. G., Starodub, D., Williams, G. J., Hajdu, J., Timneanu, N., Seibert, M. M., Andreasson, J., Rocker, A., Jonsson, O., Svenda, M., Stern, S., Nass, K., Andritschke, R., Schroter, C. D., Krasniqi, F., Bott, M., Schmidt, K. E., Wang, X., Grotjohann, I., Holton, J. M., Barends, T. R., Neutze, R., Marchesini, S., Fromme, R., Schorb, S., Rupp, D., Adolph, M., Gorkhover, T., Andersson, I., Hirsemann, H., Potdevin, G., Graafsma, H., Nilsson, B. & Spence, J. C. (2011). Nature 470, 73–77.

Chen, V. B., Arendall, W. B., 3rd, Headd, J. J., Keedy, D. A., Immormino, R. M., Kapral, G. J., Murray, L. W., Richardson, J. S. & Richardson, D. C. (2010). Acta Crystallogr D Biol Crystallogr 66, 12–21.

Emsley, P., Lohkamp, B., Scott, W. G. & Cowtan, K. (2010). Acta Crystallogr D Biol Crystallogr 66, 486–501.

Fuller, F. D., Gul, S., Chatterjee, R., Burgie, E. S., Young, I. D., Lebrette, H., Srinivas, V., Brewster, A. S., Michels-Clark, T., Clinger, J. A., Andi, B., Ibrahim, M., Pastor, E., de Lichtenberg, C., Hussein, R., Pollock, C. J., Zhang, M., Stan, C. A., Kroll, T., Fransson, T., Weninger, C., Kubin, M., Aller, P., Lassalle, L., Brauer, P., Miller, M. D., Amin, M., Koroidov, S., Roessler, C. G., Allaire, M., Sierra, R. G., Docker, P. T., Glownia, J. M., Nelson, S., Koglin, J. E., Zhu, D., Chollet, M., Song, S., Lemke, H., Liang, M., Sokaras, D., Alonso-Mori, R., Zouni, A., Messinger, J., Bergmann, U., Boal, A. K., Bollinger, J. M., Jr., Krebs, C., Hogbom, M., Phillips, G. N., Jr., Vierstra, R. D., Sauter, N. K., Orville, A. M., Kern, J., Yachandra, V. K. & Yano, J. (2017). Nat Methods 14, 443–449.

Hamilton© (2025). PipeJet® Nano Dispenser, https://www.hamiltoncompany.com/pipejet.

Henkel, A., Galchenkova, M., Maracke, J., Yefanov, O., Klopprogge, B., Hakanpaa, J., Mesters, J. R., Chapman, H. N. & Oberthuer, D. (2023). IUCrJ 10, 253–260.

Ishikawa, T., Aoyagi, H., Asaka, T., Asano, Y., Azumi, N., Bizen, T., Ego, H., Fukami, K., Fukui, T., Furukawa, Y., Goto, S., Hanaki, H., Hara, T., Hasegawa, T., Hatsui, T., Higashiya, A., Hirono, T., Hosoda, N., Ishii, M., Inagaki, T., Inubushi, Y., Itoga, T., Joti, Y., Kago, M., Kameshima, T., Kimura, H., Kirihara, Y., Kiyomichi, A., Kobayashi, T., Kondo, C., Kudo, T., Maesaka, H., Marechal, X. M., Masuda, T., Matsubara, S., Matsumoto, T., Matsushita, T., Matsui, S., Nagasono, M., Nariyama, N., Ohashi, H., Ohata, T., Ohshima, T., Ono, S., Otake, Y., Saji, C., Sakurai, T., Sato, T., Sawada, K., Seike, T., Shirasawa, K., Sugimoto, T., Suzuki, S., Takahashi, S., Takebe, H., Takeshita, K., Tamasaku, K., Tanaka, H., Tanaka, R., Tanaka, T., Togashi, T., Togawa, K., Tokuhisa, A., Tomizawa, H., Tono, K., Wu, S. K., Yabashi, M., Yamaga, M., Yamashita, A., Yanagida, K., Zhang, C., Shintake, T., Kitamura, H. & Kumagai, N. (2012). Nat Photonics 6, 540–544.

Kameshima, T., Ono, S., Kudo, T., Ozaki, K., Kirihara, Y., Kobayashi, K., Inubushi, Y., Yabashi, M., Horigome, T., Holland, A., Holland, K., Burt, D., Murao, H. & Hatsui, T. (2014). The Review of scientific instruments 85, 033110.

Kubo, M., Nango, E., Tono, K., Kimura, T., Owada, S., Song, C., Mafune, F., Miyajima, K., Takeda, Y., Kohno, J. Y., Miyauchi, N., Nakane, T., Tanaka, T., Nomura, T., Davidsson, J., Tanaka, R., Murata, M., Kameshima, T., Hatsui, T., Joti, Y., Neutze, R., Yabashi, M. & Iwata, S. (2017). J Synchrotron Radiat 24, 1086–1091.

Kumagai, I., Sunada, F., Takeda, S. & Miura, K. (1992). J Biol Chem 267, 4608–4612.

Mafune, F., Miyajima, K., Tono, K., Takeda, Y., Kohno, J. Y., Miyauchi, N., Kobayashi, J., Joti, Y., Nango, E., Iwata, S. & Yabashi, M. (2016). Acta Crystallogr D Struct Biol 72, 520–523.

McCoy, A. J., Grosse-Kunstleve, R. W., Adams, P. D., Winn, M. D., Storoni, L. C. & Read, R. J. (2007). J Appl Crystallogr 40, 658–674.

Nakane, T., Joti, Y., Tono, K., Yabashi, M., Nango, E., Iwata, S., Ishitani, R. & Nureki, O. (2016). J Appl Crystallogr 49, 1035–1041.

Nango, E., Sugahara, M., Kobayashi, J., Tanaka, T., Yamashita, A., Pan, D., Tanaka, Y., Ihara, K., Suno, C. & Shimamura, T. (2015). PSSJ Archives 8, e081.

Neutze, R., Wouts, R., van der Spoel, D., Weckert, E. & Hajdu, J. (2000). Nature 406, 752–757.

Nguyen, R. C., Yang, Y., Wang, Y., Davis, I. & Liu, A. (2020). ACS Catal 10, 1628–1639.

Roessler, C. G., Agarwal, R., Allaire, M., Alonso-Mori, R., Andi, B., Bachega, J. F. R., Bommer, M., Brewster, A. S., Browne, M. C., Chatterjee, R., Cho, E., Cohen, A. E., Cowan, M., Datwani, S., Davidson, V. L., Defever, J., Eaton, B., Ellson, R., Feng, Y., Ghislain, L. P., Glownia, J. M., Han, G., Hattne, J., Hellmich, J., Heroux, A., Ibrahim, M., Kern, J., Kuczewski, A., Lemke, H. T., Liu, P., Majlof, L., McClintock, W. M., Myers, S., Nelsen, S., Olechno, J., Orville, A. M., Sauter, N. K., Soares, A. S., Soltis, S. M., Song, H., Stearns, R. G., Tran, R., Tsai, Y., Uervirojnangkoorn, M., Wilmot, C. M., Yachandra, V., Yano, J., Yukl, E. T., Zhu, D. & Zouni, A. (2016). Structure 24, 631–640.

Schrödinger, L. & DeLano, W. (2020). http://www.pymol.org/pymol.

Shimazu, Y., Tono, K., Tanaka, T., Yamanaka, Y., Nakane, T., Mori, C., Terakado Kimura, K., Fujiwara, T., Sugahara, M., Tanaka, R., Doak, R. B., Shimamura, T., Iwata, S., Nango, E. & Yabashi, M. (2019). J Appl Crystallogr 52, 1280–1288.

Streule, W., Lindemann, T., Birkle, G., Zengerle, R. & Koltay, P. (2004). JALA: Journal of the Association for Laboratory Automation 9, 300–306.

Sugahara, M., Mizohata, E., Nango, E., Suzuki, M., Tanaka, T., Masuda, T., Tanaka, R., Shimamura, T., Tanaka, Y., Suno, C., Ihara, K., Pan, D., Kakinouchi, K., Sugiyama, S., Murata, M., Inoue, T., Tono, K., Song, C., Park, J., Kameshima, T., Hatsui, T., Joti, Y., Yabashi, M. & Iwata, S. (2015). Nat Methods 12, 61–63.

Tenboer, J., Basu, S., Zatsepin, N., Pande, K., Milathianaki, D., Frank, M., Hunter, M., Boutet, S., Williams, G. J., Koglin, J. E., Oberthuer, D., Heymann, M., Kupitz, C., Conrad, C., Coe, J., Roy-Chowdhury, S., Weierstall, U., James, D., Wang, D., Grant, T., Barty, A., Yefanov, O., Scales, J., Gati, C., Seuring, C., Srajer, V., Henning, R., Schwander, P., Fromme, R., Ourmazd, A., Moffat, K., Van Thor, J. J., Spence, J. C., Fromme, P., Chapman, H. N. & Schmidt, M. (2014). Science 346, 1242–1246.

Tono, K., Hara, T., Yabashi, M. & Tanaka, H. (2019). J Synchrotron Radiat 26, 595–602.

Tono, K., Nango, E., Sugahara, M., Song, C. Y., Park, J., Tanaka, T., Tanaka, R., Joti, Y., Kameshima, T., Ono, S., Hatsui, T., Mizohata, E., Suzuki, M., Shimamura, T., Tanaka, Y., Iwata, S. & Yabashi, M. (2015). Journal of Synchrotron Radiation 22, 532–537.

White, T. A., Barty, A., Stellato, F., Holton, J. M., Kirian, R. A., Zatsepin, N. A. & Chapman, H. N. (2013). Acta Crystallogr D Biol Crystallogr 69, 1231–1240.

Zielinski, K. A., Prester, A., Andaleeb, H., Bui, S., Yefanov, O., Catapano, L., Henkel, A., Wiedorn, M. O., Lorbeer, O., Crosas, E., Meyer, J., Mariani, V., Domaracky, M., White, T. A., Fleckenstein, H., Sarrou, I., Werner, N., Betzel, C., Rohde, H., Aepfelbacher, M., Chapman, H. N., Perbandt, M., Steiner, R. A. & Oberthuer, D. (2022). IUCrJ 9, 778–791.

