## Supporting information for "Compact Tape-Driven Sample Delivery System for Serial Femtosecond Crystallography"

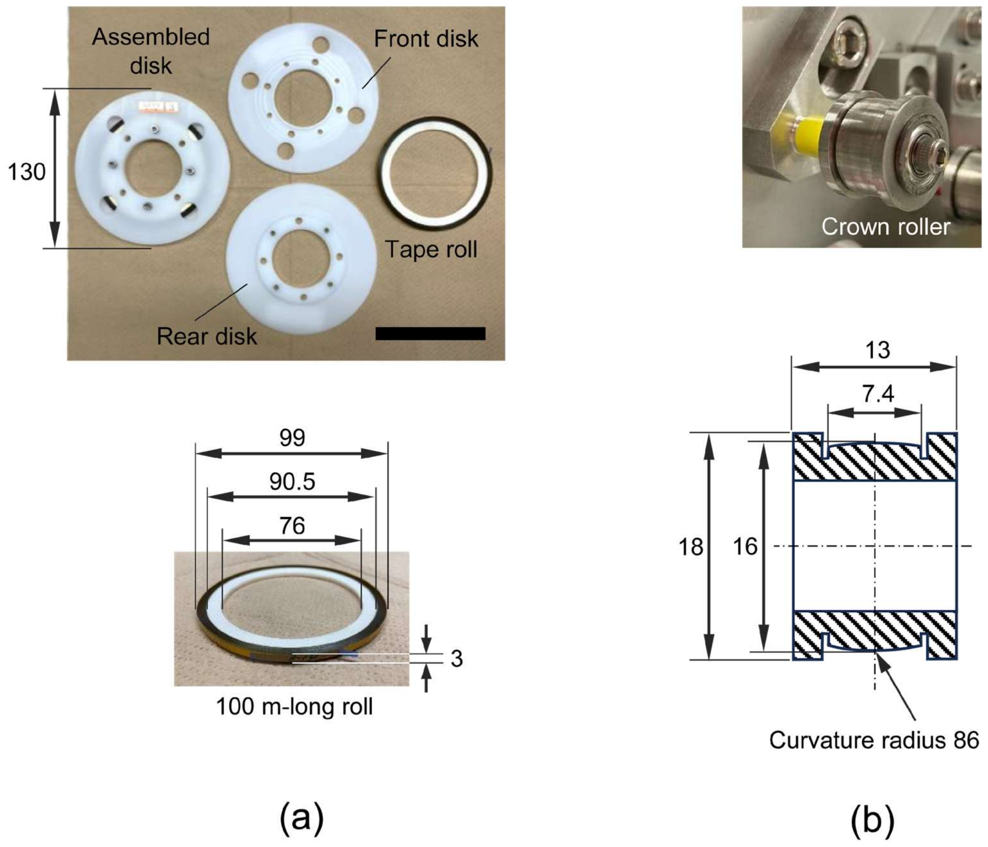


**Figure S1** Details of the CoT system components: (a) Photographs of the tape cartridge disk with a typical tape roll. The scale bar in the right-hand corner represents 100 mm. (b) Design of the crown roller. Units are in millimeters.

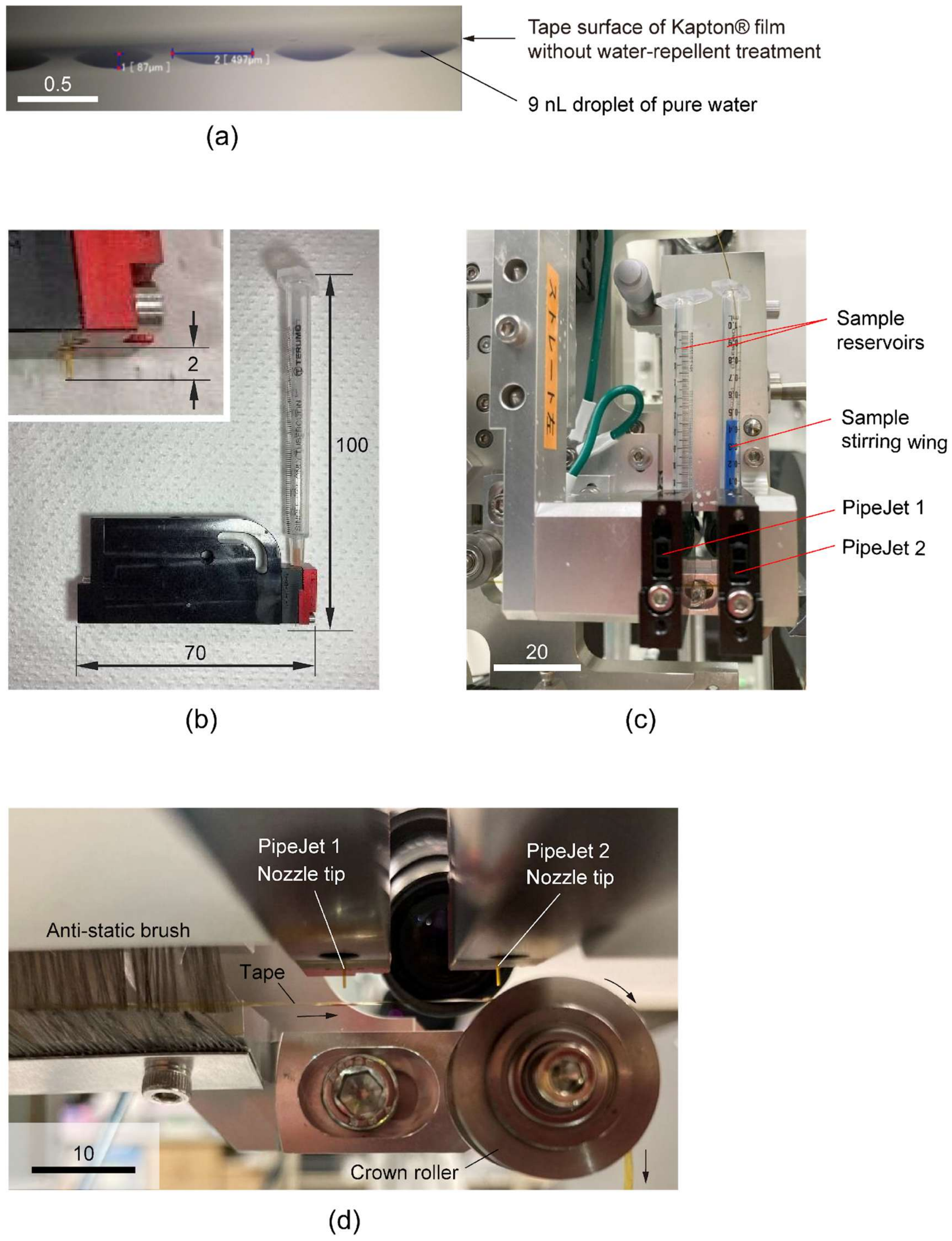


**Figure S2** Photographs of the piezoelectric injector of PipeJet® and ejected droplet queue on the tape surface: (a) 9 nL droplet queue of pure water on the tape surface of the Kapton® film without water-repellent treatment, (b) PipeJet unit mounted with a 1 mL Terumo syringe as the sample reservoir, (c) two PipeJet bodies installed on the ejection head plate with two individual sample reservoirs (PipeJet 1: substrate solution , PipeJet 2: crystal slurry) and a sample stirring wing inserted in the crystal reservoir, and (d) expanded view of the nozzle tips and their surroundings. The three arrows in (d) indicate the tape travel direction. Units are in millimeters.

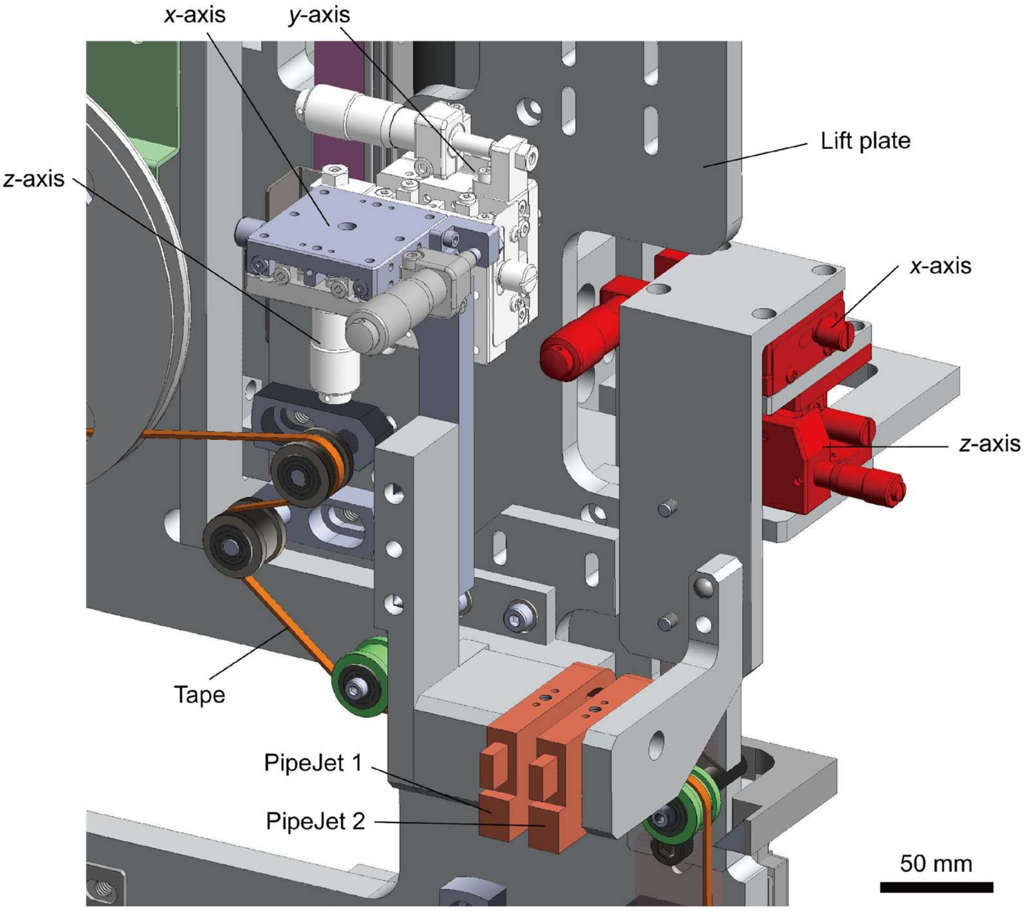


**Figure S3** CAD diagram of PipeJet® units 1 and 2 mounted on the lift plate with *x*-, *y*-, and *z*-axis micrometer stages and *x*- and *z*-axis micrometer stages, respectively.

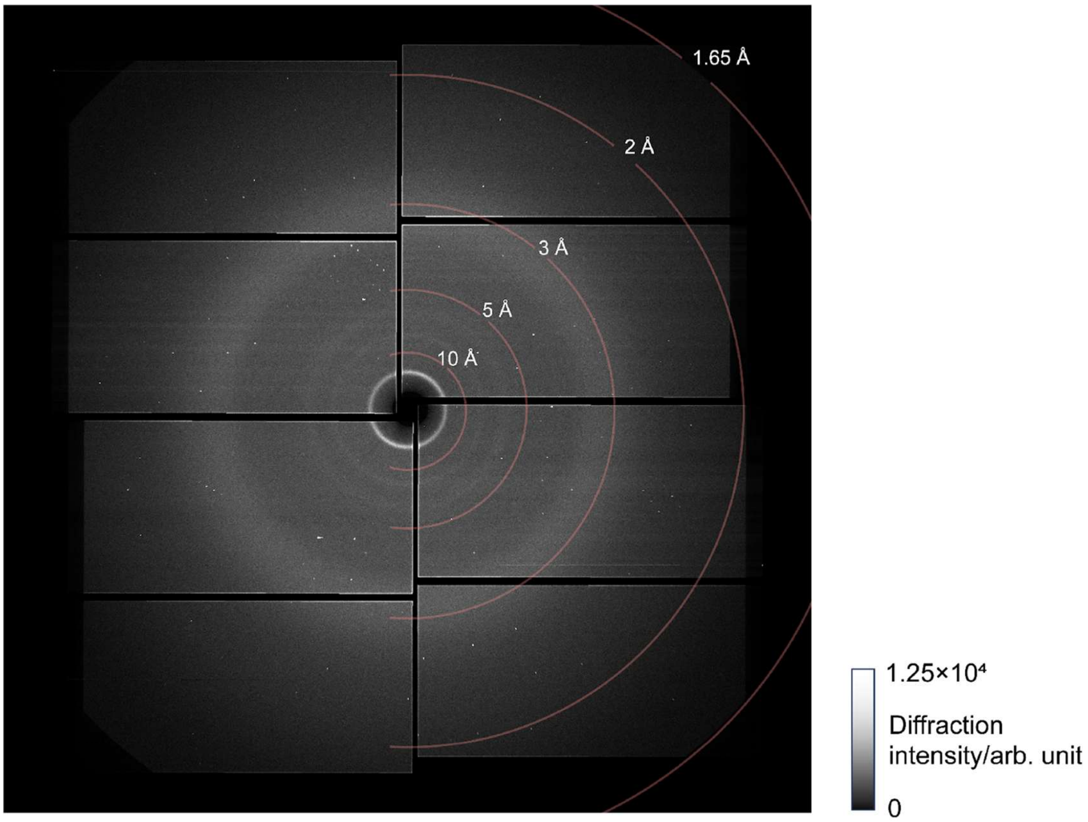
1.65 Å
2 Å
3 Å
5 Å
10 Å
1.25×10⁴
Diffraction intensity/arb. unit
0

(a)

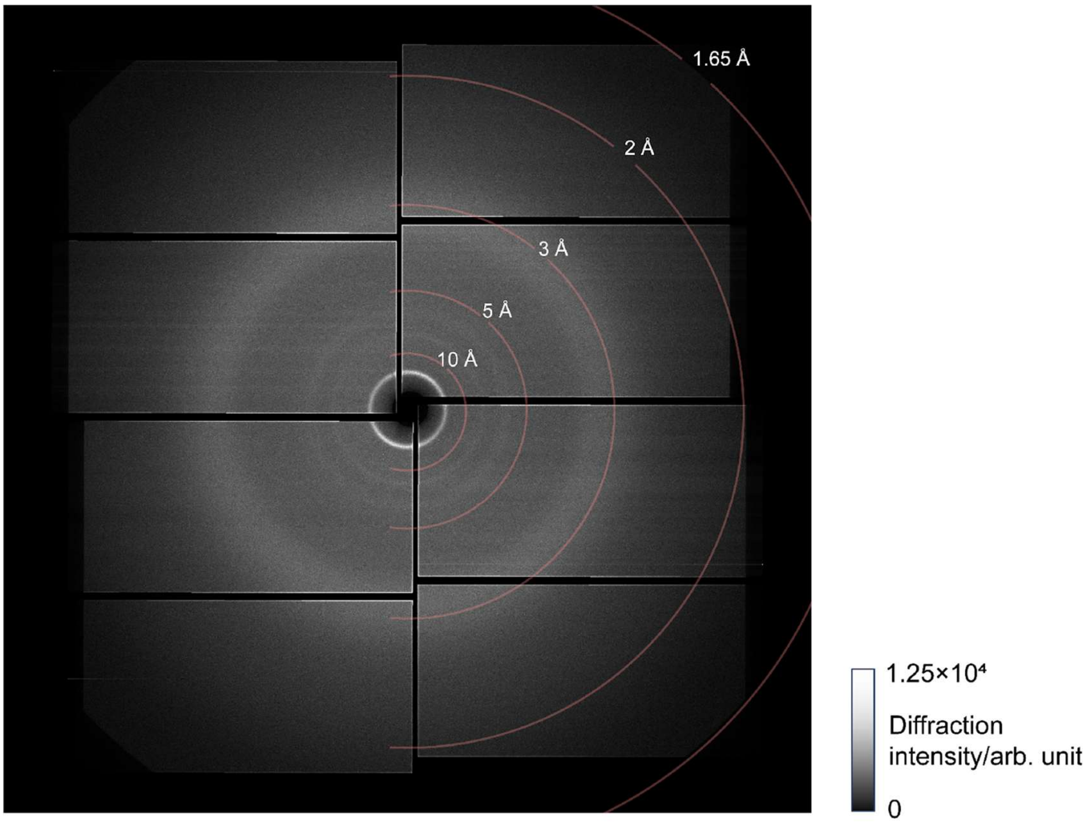
1.65 Å
2 Å
3 Å
5 Å
10 Å
1.25×10⁴
Diffraction intensity/arb. unit
0

(b)

**Figure S4** Typical XFEL diffraction patterns recorded by the phase-III MPCCD SWD. (a) The diffraction pattern of a 3–5 µm-HEWL microcrystal without GlcNAc, (b) Scattering patterns from a droplet without crystals on the tape. The XFEL pulse is focused on a 1.5 µm-diameter with the photon energy of 10 keV. The pulse energy is around 370 µJ on average. The droplet volume is from 10 to 14 nL, which is estimated to have a height of 90 to 100 µm, respectively. The distance from the sample–XFEL pulse interaction point to the detector surface is 70 mm. The white circles inside of 5 Å are diffraction rings belonging to the 12.5-µm-thick Kapton® film.

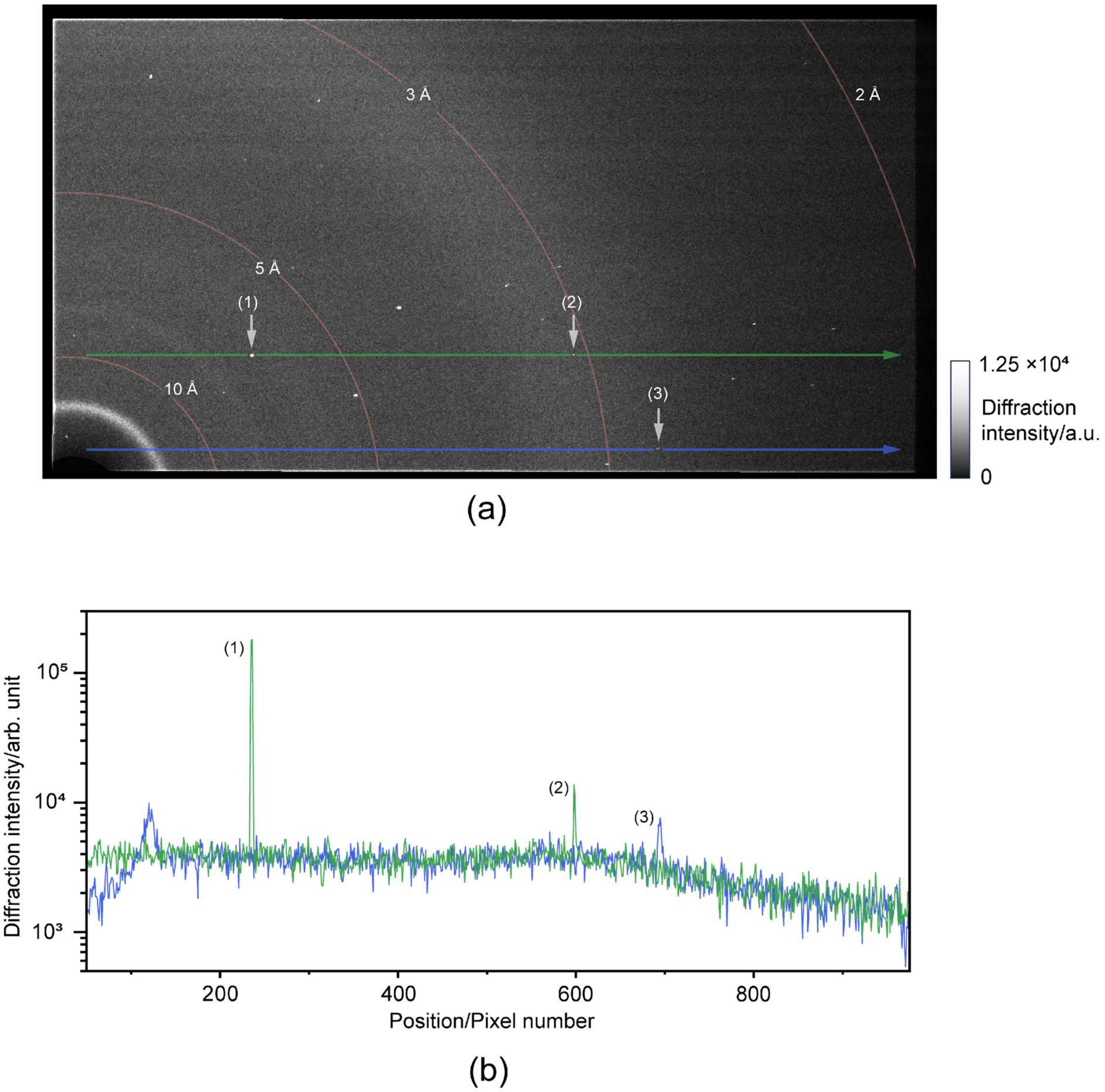


**Figure S5** (a) The magnified diffraction patterns of Fig. S4a, the second panel from the top of the right side, (b) the cross-sections of the green line arrow and blue line arrow in S5a.

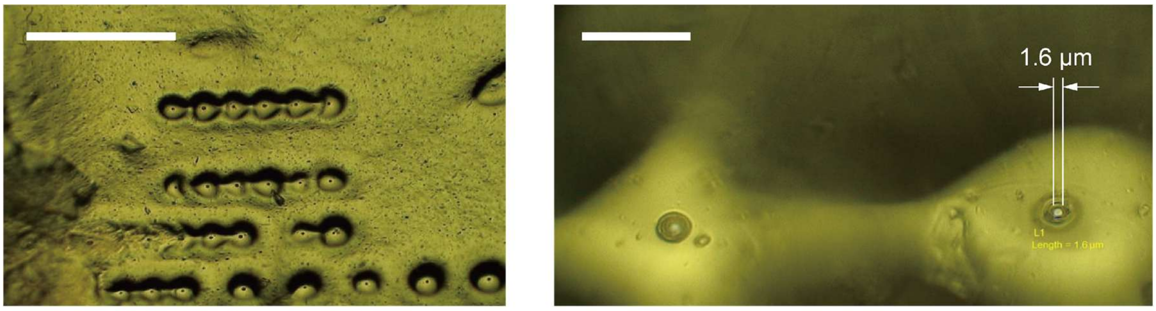


(a) (b)

**Figure S6** Photographs of the tape surface of Kapton® film after the XFEL irradiation: (a) Traces on the tape surface after XFEL irradiation, and (b) the zoomed-in view of the traces on the tape surface after the XFEL irradiation. A hole punched due to the XFEL has a diameter of 1.6 μm, which is quite close to the XFEL beam size itself. The scale bars indicate (a) 500 μm and (b) 20 μm.

**Table S1** Protein concentration estimation based on the number of molecules in HEWL crystals

| Crystal size | 1-µm HEWL crystal | 3–5-µm HEWL crystal |
|---|---|---|
| Protein molecular weight | 14,296 Da (2.4×$10^{-17}$ mg) | |
| Protein molecular number per units | 8 | |
| Unit cell volume | 2.4×$10^{-7}$ µ$m^3$ (2.4×$10^{-13}$ nL) | |
| Crystal volume | 1 µ$m^3$ (1×$10^{-6}$ nL) | 1.2×$10^{2}$ µ$m^3$ (1.2×$10^{-4}$ nL) |
| Unit cell number per crystal | 4.2×$10^{6}$ | 4.9×$10^{8}$ |
| Protein molecule number per crystal | 3.3×$10^{7}$ | 3.9×$10^{9}$ |
| Protein mass per crystal | 7.9×$10^{-10}$ mg | 9.3×$10^{-8}$ mg |
| Crystal density | 1.1×$10^{9}$ crystals m$L^{-1}$ | 4.8×$10^{8}$ crystals m$L^{-1}$ |
| Protein concentration | 8.7×$10^{-1}$ mg m$L^{-1}$ | 4.4×$10^{1}$ mg m$L^{-1}$ |

**Table S2** Number of crystals and protein mass in a droplet

| Droplet volume | 10 nL (crystal slurry) | 14 nL (crystal slurry) | 20 nL (combined 10 nL droplet of crystal slurry & 10 nL droplet of inhibitor solution) | 28 nL (combined 14 nL droplet of crystal slurry & 14 nL droplet of inhibitor solution) |
|---|---|---|---|---|
| Diameter (µm) | 520 | 580 | 660 | 740 |
| Height (µm) | 91 | 102 | 116 | 130 |
| 1 µm crystal numbers in a droplet (1.1×$10^{9}$ crystals/mL) | 11,000 | 15,400 | 11,000 | 15,400 |
| 3–5 µm crystal numbers in a droplet (4.8×$10^{8}$ crystals/mL) | 4,800 | 6,720 | 4,800 | 6,720 |
| Protein mass per droplet for 1 µm crystal (µg) | 8.7×$10^{-3}$ | 1.2×$10^{-2}$ | 8.7×$10^{-3}$ | 1.2×$10^{-2}$ |
| Protein mass per droplet for 3–5 µm crystal (µg) | 4.4×$10^{-1}$ | 6.2×$10^{-1}$ | 4.4×$10^{-1}$ | 6.2×$10^{-1}$ |
| Total volume ratio of 1 µm crystal per droplet volume (%) | 0.1 | | | |
| Total volume ratio of 3–5 µm crystal per droplet volume (%) | 5.7 | | | |
| 1 µm crystal numbers in the 1.5 µm-diameter XFEL pulse path | 0.18 | 0.20 | 0.11 | 0.13 |
| 3–5 µm crystal numbers in the 1.5 µm-diameter XFEL pulse path | 0.08 | 0.09 | 0.05 | 0.06 |

**Table S3** Sample consumption per dataset

| Data set | Elapsed measurement time (second) | Indexed image | Total consumed protein weight per data set (mg) / Consumed protein weight per 10k indexed image (mg) | | |
|---|---|---|---|---|---|
| | | | **10 nL droplet** | **14 nL droplet** | **Average** |
| 3–5 µm crystal without inhibitor | 1,070 | 12,213 | 14.1 / 11.6 | 19.9 / 16.3 | 17.0 / 13.9 |
| 3–5 µm crystal 226 mM GlcNAc 2 s | 890 | 10,062 | 11.7 / 11.7 | 16.6 / 16.5 | 14.2 / 14.1 |
| 3–5 µm crystal 226 mM GlcNAc 5 s | 1,958 | 20,382 | 25.8 / 12.7 | 36.4 / 17.9 | 31.1 / 15.3 |
| 3–5 µm crystal 226 mM GlcNAc 9.7 s | 1,810 | 16,607 | 23.9 / 14.9 | 33.7 / 21.0 | 28.8 / 17.9 |
| 1 µm crystal without inhibitor | 3,354 | 23,501 | 0.9 / 0.4 | 1.2 / 0.5 | 1.0 / 0.4 |
| 1 µm crystal 226 mM GlcNAc 1.3 s | 6,632 | 39,020 | 1.7 / 0.4 | 2.4 / 0.6 | 2.1 / 0.5 |
| 1 µm crystal 226 mM GlcNAc 5 s | 3,860 | 16,967 | 1.1 / 0.6 | 1.4 / 0.8 | 1.2 / 0.7 |
| 1 µm crystal 226 mM GlcNAc 7.5 s | 2,136 | 11,967 | 0.6 / 0.5 | 0.8 / 0.6 | 0.7 / 0.6 |
| 1 µm crystal 226 mM GlcNAc 9.7 s | 5,344 | 17,407 | 1.4 / 0.8 | 1.9 / 1.1 | 1.7 / 1.0 |
| 1 µm crystal 452 mM GlcNAc 1.3 s | 2,563 | 15,331 | 0.7 / 0.4 | 0.9 / 0.6 | 0.8 / 0.5 |
| 1 µm crystal 452 mM GlcNAc 2.5 s | 2,139 | 12,483 | 0.6 / 0.5 | 0.8 / 0.6 | 0.7 / 0.5 |
| 1 µm crystal 452 mM GlcNAc 4 s | 3,070 | 12,723 | 0.8 / 0.6 | 1.1 / 0.9 | 1.0 / 0.8 |
| 1 µm crystal 452 mM GlcNAc 5 s | 2,351 | 17,626 | 0.6 / 0.4 | 0.9 / 0.5 | 0.7 / 0.4 |
